# Charting the Spatial Transcriptome of the Human Cerebral Cortex at Single-Cell Resolution

**DOI:** 10.1101/2024.01.31.576150

**Authors:** Songren Wei, Meng Luo, Pingping Wang, Rui Chen, Xiyun Jin, Chang Xu, Xiaoyu Lin, Zhaochun Xu, Hongxin Liu, Peng Huang, Jiangping Xu, Qinghua Jiang

**Affiliations:** School of Interdisciplinary Medicine and Engineering, Harbin Medical University, Harbin 150000, China; Department of Neuropharmacology, Southern Medical University, Guangzhou 510515, China; Center for Brain Science and Brain-Inspired Intelligence, Guangdong-Hong Kong-Macao Greater Bay Area, Guangzhou 510515, China; School of Life Science and Technology, Harbin Institute of Technology, Harbin 150000, China; Department of Forensic Medicine, Guangdong Medical University, Dongguan 523808, China; Key Laboratory of Mental Health of the Ministry of Education, Southern Medical University, Guangzhou 510515, China; Foshan Maternity & Child Healthcare Hospital, Southern Medical University, Foshan 528000, China

## Abstract

In our pursuit of creating a comprehensive human cortical atlas to understand human intelligence, we examined the single-nuclei transcriptomes of 307,738 cells alongside spatial transcriptomics data from 46,948 VISIUM spots and 1,355,582 Stereo cells. Atlases reveal distinct expression patterns and spatial arrangements of cortical neural cell types. Glutamatergic neurons exhibit precise laminar patterns, often mirroring expression patterns in adjacent cortical regions. Overlaying our atlas with functional networks delineated substantial correlations between neural cell types and cortical region function. Notably, regions involved in processing sensory information (pain) display a pronounced accumulation of extratelencephalic neurons. Additionally, our atlas enabled precise localization of the thicker layer 4 of the visual cortex and an in-depth study of the stabilize the subplate structure, known as layer 6b, revealed specific marker genes and cellular compositions. Collectively, our research sheds light on the cellular foundations of the intricate and intelligent regions within the human cortex.

## Main

Diverse high cognitive functions were supported by cellular circuits formed by billion neural cells in different cerebral cortex regions^1–3^. High-resolution “cytoarchitectural atlas” of the human cerebral cortex is pivotal for understanding the mechanism of the cortical region performing specific functions. Substantial application of many recent techniques has revealed the high diversity of neuronal cells, providing an unprecedented opportunity for the unbiased exploration of neural cell types^3–11^.

In this study, we built a comprehensive cytoarchitectural atlas of the adult human cerebral cortex. We characterized the panorama of the cellular diversity and spatial structure of the human cortex by snRNA-seq and spatial transcriptomics (ST) across multiple regions. The integration of these two omics uncovers exceedingly diverse cell types and spatial arrangements in the human cortex. Meanwhile, to explore the correlation between the cellular architecture of various cortical regions and their associated functions, we integrated datasets of cortical region function assessed by functional magnetic resonance imaging (fMRI) with our transcriptional cortical atlase^12^. We delineated the cell-type function preference by calculating the correlation between multiple region functions and cell-type specificity, providing references for the neural cellular basis of functional specificity of cortical regions. Besides, we evaluated neurological disease susceptibility across cortical regions and cell types based on the neurological risk gene dataset to outline a global landscape of susceptibility at fine cell type and spatial level. The cellular atlas of the adult human cortex also empowers us to locate layer 4 and search for finer cortical structures, layer 6b, as well as discover their molecular features and highly relevant cell types. Together, our comprehensive atlas of cellular diversity and spatial arrangement in the human cortex provided the molecular and cellular basis of human intelligence.

## Results

### Transcriptomic cell-type taxonomy in the human cerebral cortex

To investigate the cellular diversity across the human cerebral cortex, snRNA-seq was performed on 42 cerebral cortical samples from 5 adult human donors without neurological disorders (Fig.1a), which covers 14 cortical regions: frontopolar prefrontal cortex (FPPFC), dorsolateral prefrontal cortex (DLPFC), ventrolateral prefrontal cortex (VLPFC), primary motor cortex (M1), anterior cingulate cortex (ACC), primary somatosensory cortex (S1), primary somatosensory cortex of the eardrum (S1E), superior temporal gyrus (STG), inferior temporal gyrus (ITG), postcentral gyrus (PoCG), supramarginal gyrus (SMG), superior parietal lobule (SPL), angular gyrus (AG) and primary visual cortex (V1) (Fig.1b, Supplementary Table 1).

**Fig. 1.**
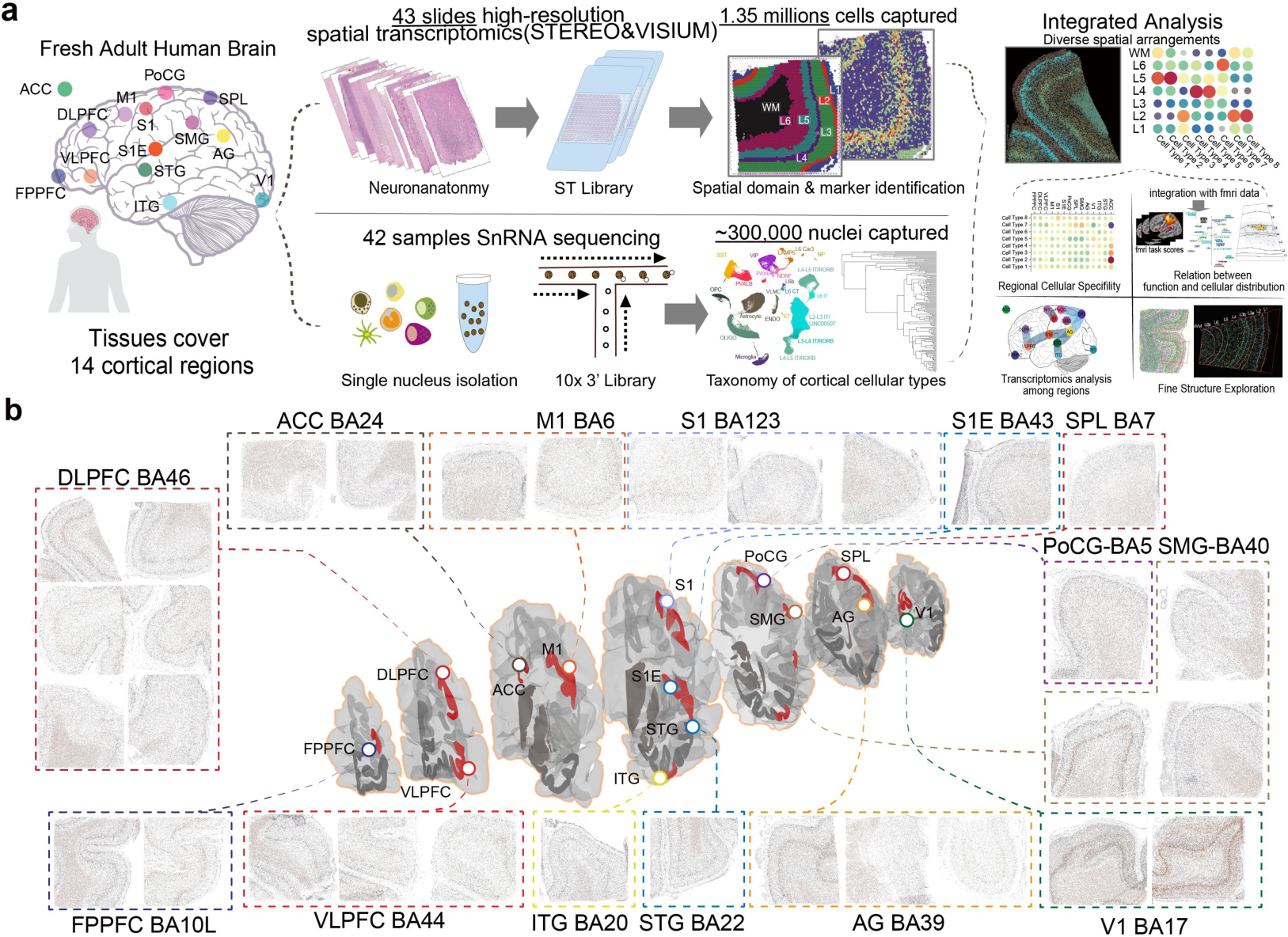
Comprehensive spatial transcriptomic atlas of the 14 human cortical regions. **a** Overview of sampled cortical regions rendered in coronal sections and schematic workflow of snRNA and ST sequencing and analysis. We obtain a comprehensive neural cell census and localize various cell types by the 10X Genomics Chromium snRNA-seq, VISIUM, and Stereo spatial transcriptomics (ST) technologies. **b** The coronal slides cover the following 14 cortical regions by using Stereo-seq. Colors denoted different coronal cortical sections, and the red color shows the part for dissection same with Extended Data Fig. 1a.

Glutamatergic and GABAergic neurons, predominantly located in the cortex, serve as fundamental components in neural circuits by primarily exerting excitatory and inhibitory influences, respectively, within the cortical neural network. Identified transcriptionally, GABAergic neurons exhibit the expression of glutamate decarboxylase (either *GAD1* or *GAD2*), which is pivotal for synthesizing the inhibitory neurotransmitter GABA^13^. Conversely, cells that express *SLC17A7*, a vesicular glutamate transporter, are typically categorized as glutamatergic neurons due to their capability to release the excitatory neurotransmitter, glutamate^4^. While subclasses of GABAergic and glutamatergic neurons can possess distinctive transcriptional profiles, morphologies, and functions, the interplay and relationships between these classifications can be complex, as underscored by the notable diversity within GABAergic and glutamatergic neuronal populations, as demonstrated by recent research ^14–16^. Our atlas, defined by transcriptomic molecular features, generally asserts that glutamatergic and GABAergic neurons converge as overlapping inhibitory/excitatory neurons. We sequenced 307,738 nuclei with a median of 3,928 genes (Supplementary Table 1). We adopted the recursive clustering pipeline^4,10,14^ for cellular clustering and defined 242 transcriptomically distinct neural cell clusters belonging to 21 subclasses under 3 major classes (Extended Data Fig. 1, and Supplementary Tables 2): 9 subclasses and 83 clusters under glutamatergic neuron class, 6 subclasses and 124 clusters under GABAergic neuron class, and 6 subclasses and 38 clusters under non-neuronal cell class (Extended Data Fig. 1). Each cell type distributed evenly in each donor, indicating the uniformity of sampling in each cortical region. Given that oligodendrocyte precursor cells and GABAergic neurons could both be produced by progenitors in the ventral telencephalon^17^, these original similarities between these two cells correspond to adjacent spatial relation in the expression (Extended Data Fig. 1). More cell types were defined within the GABAergic and non-neuronal classes in the broad coverage of cortical regions (Extended Data Fig. 1). Cajal–Retzius cells were not found due to a ratio of less than 0.1% of L1 neurons in the human cortex^18^.

### Laminar structures in the human cerebral cortex revealed by spatial transcriptomics

To delineate the cortical spatial heterogeneity, spatial transcriptome sequencing was performed on 32 Stereo slides of 14 cortical regions about ∼100k bins/slide, and with median cell number of 43461±13271/slide (Supplementary Table 1). To identify robust laminar structure, we adopted a graph-based deep learning framework^19^ to integrate multimodal data like histology (HE and ssDNA staining) and gene expression and achieved robust spatial domain identification (Fig. 2a and Extended Data Fig. 2a).

**Fig. 2.**
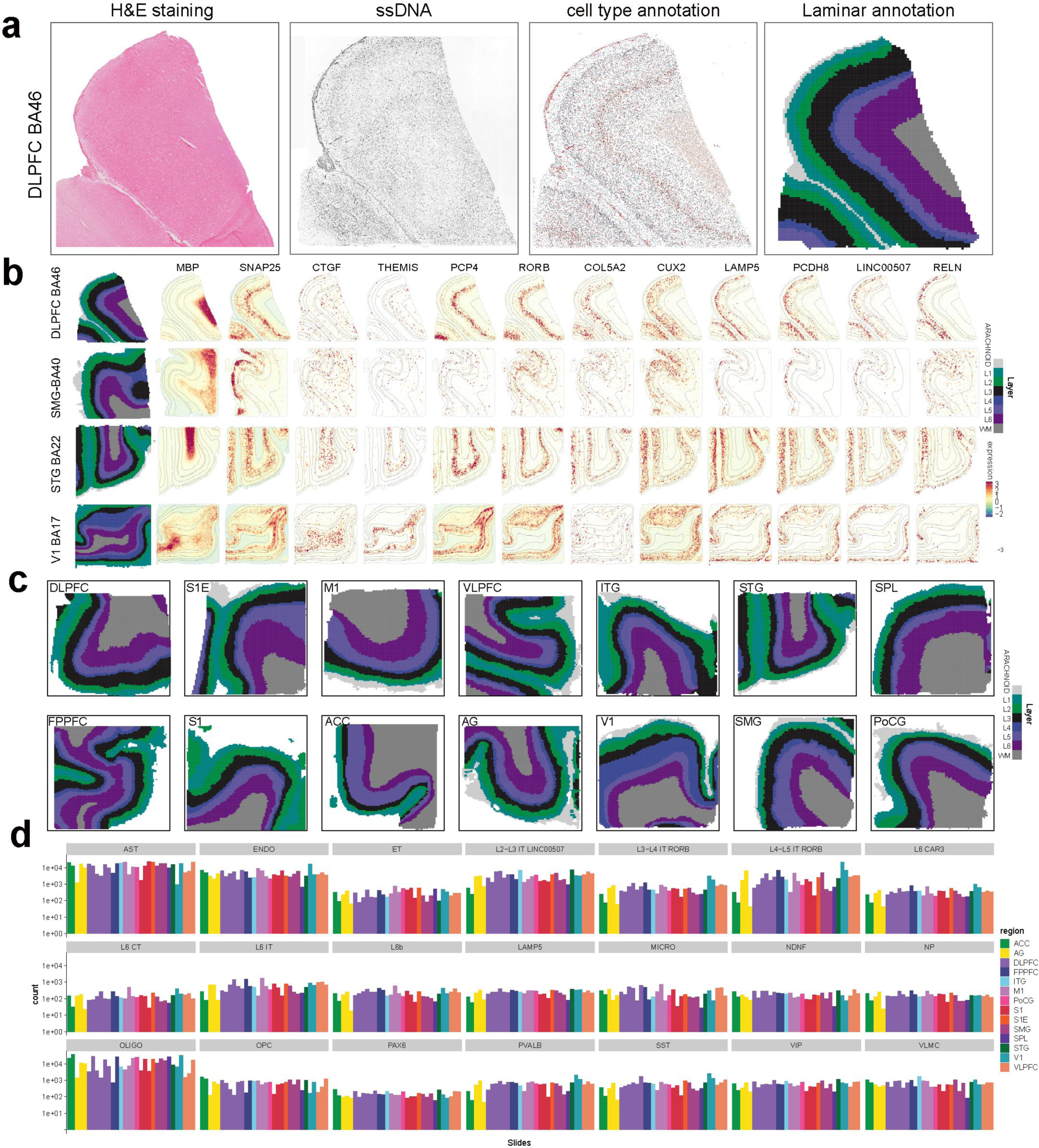
Spatial transcriptomics of cortical regions of humans and layer enrichment of previously identified layer-marker genes at Stereo-seq. **a** Single-cell and laminar segmentation for Stereo-seq analysis. H&E staining was performed on the previous section, mainly to observe the morphological structure inside the cells; (ssDNA) Total mRNA captured from an DLPFC section; (cell-type annotation) Subclass neuronal cell types was determined by clustering and RNA expression; (Laminar annotation) DeepST AI-assisted semi-supervised laminar segmentation based on nucleic acid and cell-type annotation (Methods). A gradient color scheme is employed to distinctly showcase the different layers within each cortical region. The slide size is 1×1cm. Each slide has completed transcriptomic segmentation mostly with L1 to WM. We captured a total of 1,355,582 cells from 32 slides. **b** Spotplots of normalized expression for sample DLPFC, SMG, STG and V1 for genes MBP (marker for WM), SNAP25 (marker for Neuron), CCN2 (marker for layer 6b), THEMIS (marker for layer 6), PCP4 (marker for around layer 5), RORB (marker for around layer 4), COL5A2 (marker for layer 4 and layer 3), CUX2 (marker for layer 3 and layer 2), LAMP5 (marker for layer 2), PCDH8 (marker for layer 2), LINC00507 (marker for layer 2) and RELN (marker for layer 1). **c** The remaining 13 cortical area for laminar annotation, and the median gene number about 516. **d** The statistics of cell number of different subclasses in Stereo slides, color-coded based on the region.

Various layers differed dramatically in both expression patterns and layer-specific cell types (Extended Data Fig. 2). The transcriptomic expression of spots alters with spatial position, characterizing the spatial transcriptomic continuity in the cortex, consistent with prevalent neurobiological views (Extended Data Fig. 2b-e)^20,21^. A series of laminar markers can accurately locate different layers and showed expression contiguity along with laminar changing (Fig. 2b, Extended Data Fig. 2f, and Supplementary Table 3): CCN2 highlights layer 6 and areas near WM, PCP4 highlights layer 5 and the areas near layer 4, CUX2 and COL5A2 for layer 3, LAMP5 for layer 2, LINC00507/PCDH8 for layer 2, and RELN for layer 1. Classic markers such as MBP and THEMIS are the top markers for WM and layer 6. The supragranular layers are identified combinatorial by the markers RORB and dense cellular layer staining. (Fig. 2b, and Extended Data Fig. 2f-g). These markers display exceedingly restricted to specific laminar (Fig. 2c, Extended Data Fig. 2h-j), indicating satisfactory conserveness for precise laminar markers. At the same time, we can calculate the proportion of each subclasses in each Stereo slide and find that the proportion of superficial L2-L3 IT in the anterior cingulate gyrus (ACC) is low, while other subclasses are relatively evenly distributed in various cortical regions (Fig. 2d).

### Prevalent unimodal laminar distributions of glutamatergic neurons

Glutamatergic neurons are characterized by diverse projection patterns through which axons (long, branching processes that transmit electrical signals) send the neurotransmitter glutamate to other brain regions^22^. These projection patterns imply the specific brain regions they connect to and the functions they perform in those regions, so subclasses of glutamatergic neurons can be subdivided by projection patterns^23–26^. We leveraged the single-cell transcriptome data to divide glutamatergic neurons into 83 transcriptomically distinct clusters under 9 subclasses (Fig. 3a-b, and Extended Data Fig. 3). We utilized Stereo slides to cover 14 cortical regions to observed that intratelencephalic (IT) neurons formed a continuous spectrum of cells with highly correlated gradual changes in gene expression and spatial position (Fig. 3c). Most glutamatergic clusters present unimodal laminar distribution, with a single laminar center with the highest cellular density extending to adjacent layers gradient (Extended Data Fig. 4). Detailed, IT includes 4 subclasses: the IT LINC00507 subclass in the supragranular, the L3-L4 and L4-L5 IT RORB subclasses bridging the supra- and subgranular layers, and the IT L6 subclass beneath layer 5 (Fig. 3c, d). Other subclasses correspond to different projection types, including the extratelencephalic (ET), near-projecting (NP), L6 Car3, corticothalamic (CT), and layer 6b (L6b) (Extended Data Fig. 4). Extratelencephalic (ET), proximal projection (NP), and corticothalamic (CT) neurons embody distinct classes of projection neurons, each presenting unique roles and projection patterns within the central nervous system. To unravel their relationship with the transcriptome, it is pivotal to delve into the gene expression characteristic of each neuron type. ET neurons, known for their long-term projection and integration functions, are anticipated to exhibit a transcriptome that reflects these capabilities. Specifically, genes pertinent to axonal maintenance, long-distance signal transmission, and synaptic connections to distant targets might be upregulated in ET neurons, prominently expressing transcription factors like *FEZF2*, *CTIP2*, and *BCL6*, predominantly in layer 5^27,28^. It’s noteworthy that *FEZF2* is not exclusive to ET neurons; it also finds expression in near-projecting pyramidal neurons. The NP neurons, on the other hand, might exhibit a transcriptome supporting local circuitry and short-distance axonal transport, facilitating the fine-tuning of synaptic transmission within local networks. Consequently, a distinct set of genes like *SYT6*, responsible for controlling short-range communication and potentially ensuring high fidelity or specificity in synaptic connections, might be activated^29^. Lastly, CT neurons may be characterized by the expression of a unique gene subset, including the transcription factor marker *FOXP2*, with specificity often observed in layer 6, enabling them to maintain long axons and forge precise connections within the thalamus^7^.

**Fig. 3.**
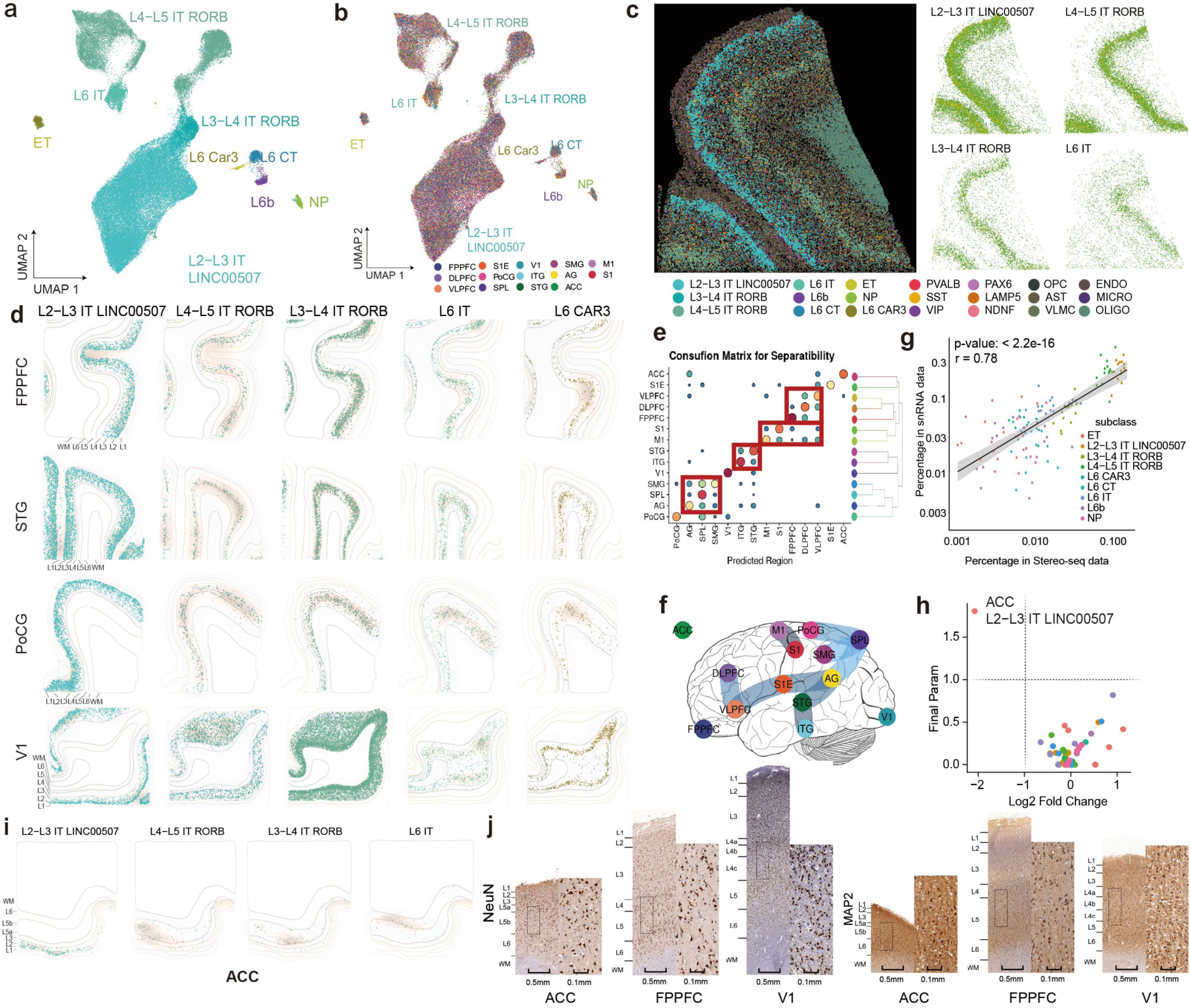
Transcriptomic taxonomy of glutamatergic cell types and spatial arrangement in the human cerebral cortex. **a** The UMAP projection plot of glutamatergic neurons. glutamatergic neurons were divided into 9 subclasses according to their projection types: NP, L6 car3, L6 IT, ET, L6 CT, L6B, L2-L3/IT, L3-L4/IT and L4-L5/IT. **b** Detailed UMAP plot showcasing diverse glutamatergic subclasses and their regional relationships. **c** The spatially resolved single-cell transcriptome of the adult human telencephalon DLPFC as determined by Stereo-seq analyses (left). Scatterplot of Stereo-seq example (DLPFC) showing 4 gradient-laminar distributed glutamatergic subclasses (right). Cells are colored by their subclass ID (same color as in Extended Data Fig. 1a). The slide size is 1×1cm. **d** Laminar distribution of typical glutamatergic neurons at the Stereo spatial cortical regions. Dot plots show the Stereo spatial transcriptome results of the glutamatergic neurons divided into 5 different subclasses. The color and alpha of spots denote the position of each neuron. **e, f** Regional transcriptomic similarities between cortical regions. Heatmap of the confusion matrix showing the separability of glutamatergic neurons from different cortical regions (**e**). Constellation plots of regional transcriptomic similarities between cortical regions. The link between the two regions delineated the shared cell proportion in K-nearest neighbors (KNN) discrimination with the region as the label and transcriptome as the feature (**f**). Only the link with shared fraction over 5% was shown. **g** Scatterplot depicting the percentages of various glutamatergic subclasses in two datasets (snRNA-seq and Stereo-seq), showcasing the correlation of Stereo-seq data with previously established findings. Each dot denotes a subclass of cells in region whose x axis coordination is its percentage in stereo datasets and y axis coordination is its percentage in snRNA-seq. These two datasets show a high consistency in cellular compositions. **h** The cellular composition analysis show differential subclass in snRNA-seq datasets, highlighting the variability of LINC00507 IT neurons in the ACC. Each dot denotes cells in a subclass in one region compared to all others regions. **i** Scatterplot of Stereo-seq example (ACC) showing 4 gradient-laminar distributed glutamatergic IT subclasses. Cells are colored by their cell density. The slide size is 1cm x 1cm. **j** Mature neurons (NeuN) and projection neurons (MAP2) distribution. Left, NeuN and MAP2 staining for neuron. Right, laminar distribution of mature and projection neurons under immunohistochemical staining.

To further investigate cross-regional variation, we evaluated the similarity of these regions based on the transcriptomic separability of glutamatergic neurons and observed the specific and continuous variability of glutamatergic neuron subclasses across cortical regions (Fig. 3e, f, and Extended Data Fig. 5-6). In almost all cases, the glutamatergic subclasses contributed similarly to adjacent cortical regions, especially in the prefrontal triangle (FPPFC, DLPFC, and VLPFC) and the angular gyrus-connecting parietal triangle (AG, SMG, and SPL) (Fig. 3d, e). We observed a high degree of consistency between the proportions of excitatory neuron subclasses in single-cell data and the Stereo-slides (Fig. 3f), suggesting a robust correlation between the two datasets. We found that the proportion of L2-L3 IT in the anterior cingulate cortex (ACC) lower in the single-cell data (Fig. 3g), and the Stereo slide displays also confirmed this trend (Fig. 3h).

The analyzed areas exhibit clear differences in cytoarchitecture, as evidenced by immunohistochemical staining using markers such as NeuN, and MAP2. This staining highlights variations in cell size, shape, laminar and columnar organization, all of which extend across the rostrocaudal and mediolateral axes of the cortical sheet (Extended Data Fig. 7). In contrast to V1, which features an expanded and highly specialized layer 4, the ACC does not possess a layer 4 and exhibits a lower proportion of superficial IT neurons compared to other regions. Comprehensive observations through immunohistochemistry and Stereo slides unveiled the presence of ET (Von Economo neurons) within the ACC. Notably, both layer 5 and layer 6 in ACC occupy significant portions of the gray matter area, thereby compressing the supragranular layer that is typically populated by superficial neurons. (Fig. 3i).

Meanwhile, diffusion weighted imaging and fiber tractography provide a unique noninvasive technique to study the macro connectivity structure within the brain^30^. By quantifying such data, we generate a connectome matrix which reveals that higher order functions like working memory score highly in the prefrontal lobe, while primary sensory functions (motor and pain back) score higher in the central sulcus (Extended Data Fig. 8). Further linkage with our cellular atlas suggests that the higher order functions of the prefrontal lobe may be related to the higher density of RORB cell types within this cortical region, while the primary sensory functions of the central sulcus may be more relevant to ET (Fig. 4a). We found that the pain-related cortical regions in the cortical functional imaging were adjacent to the postcentral sulcus (S1, S1E, PoCG) and anterior cingulate (ACC) (Fig. 4b, and Extended Data Fig. 8). We also found that ET in the postcentral sulcus is densely located in layer 5 (Fig. 4b). It is speculated that the dense appearance of ET in layer 5 in the neocortex is highly related to the pain-related functions of the cortical region. However, the ACC belongs to the anterior neocortex and is not functionally related to ET neurons in pain-related cortical regions. Von Economo neurons (VEN), large spindle-shaped cells, are expected to be ET in the human telencephalon, and they are restricted to the ACC and frontal insula (FI)^27^. We also found that these VEN marker genes described as FI region did not apply to ET in the ACC region, and there were only two ET clusters in the ACC, which were generally highly expressed *RREB1*, *TEC*, *ADAMTS2*, and *ITGA4* were not expressed (Fig. 4c). Gene ontology terms were performed for ET in the ACC, and they were found to be associated with potassium ion transmembrane transporter activity (Fig. 4d). KEGG and GO term analysis also found substantial differences on other subclasses (Extended Data Fig. 9-10), and disease susceptibility analysis was performed across multiple cortical regions and subclasses of glutamatergic neurons by overlaying them with neural risk gene sets (Extended Data Fig. 11).

**Fig. 4.**
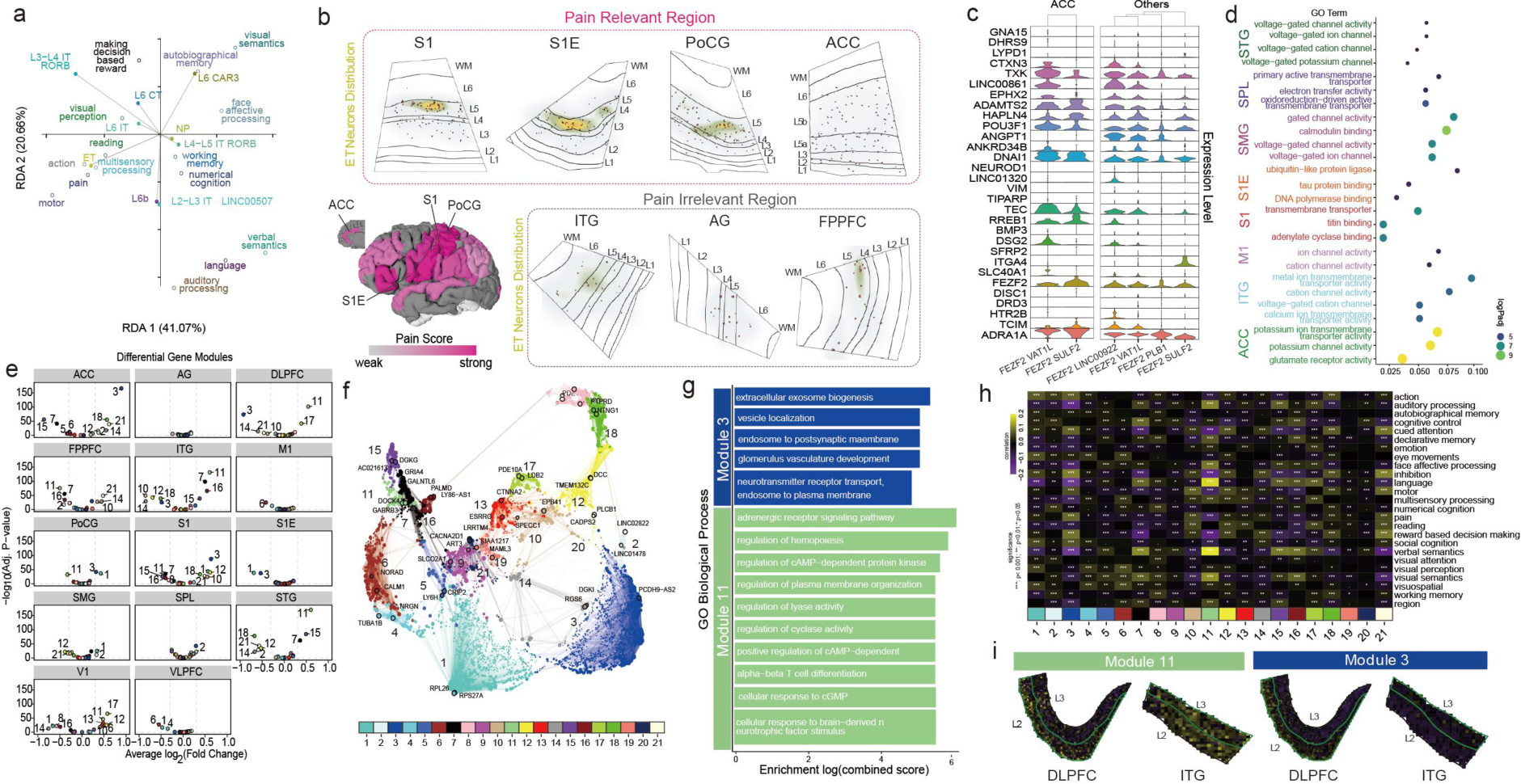
Functional enrichment in pain-relevant and irrelevant regions of the human cortex, and WGCNA-weighted gene co-expression. **a** The biplot showed the correlation between cortical region function and cell specificity by redundancy analysis (RDA). Only the top 2 nearest subclasses to each topic were marked. The angle reflects the correlation between function and cell type, and the length indicates the magnitude of each explicative contribution. **b** Detailed representation of specific cortical regions relevant to pain perception. Zoom-in Stereo slides view for pain-relevant (S1E, PoCG, ACC) and pain-irrelevant (ITG, AG, FPPFC) cortical regions, illustrating laminar organization and the distribution of ET. ET densely distributed in pain relevant regions such as S1, S1E, PoCG, while the region with least correlation with pain were found much less ET. The down left plot denotes the pain correlation in cortical regions. **c** Violin plots in ACC (left) and others (right) show the expression of markers in ET. **d** Dot plot representing the GO term enrichment analysis for specific regions, highlighting the molecular functions and biological processes of ET in multi-cortical regions. **e** Volcano plot shows differential expression of 21 gene co-expression modules in L2-L3 IT in different cortical regions. **f** UMAP plot of genes show hub genes in each gene module. The UMAP is calculated by the similarity of expression in cells and colored by gene modules. **g** The pathways enriched by gene modules 3 and 11 from L2-L3 IT. **h** The trait test show the correlation between L2-L3 IT genes modules and different cortical functions. **i** The expression of gene module 11 and module 3 in region DLPFC and ITG.

WGCNA-weighted gene co-expression module network analysis was performed on 14 cortical regions using single-cell data, and co-expression gene modules of multiple subclasses were explored (Fig. 4e-g). It was found that in the L2-L3 IT LINC00507, a 3-gene module was highly expressed in ACC and S1, and low expression in DLPFC and ITG (Fig. 4e). The 3-gene module is related to kinetochore microtubule and extracellular vesicles in biological processes (Fig. 4g). At the same time, it was found that an 11-module was highly expressed in DLPFC and ITG, and this module is related to cAMP-dependent regulatory pathways (Fig. 4g). Correlation analysis between gene modules and task scores in FMRI cortical regions found that module 11 is highly correlated with language and speech semantic tasks (Fig. 4h), indicating that language-related scores require more functions of module 11 in L2-L3 IT. Meanwhile, we also observed through Stere-seq that module 11 is highly expressed in the superficial layer 2-3 in DLPFC and ITG, while module 3 is low-expressed (Fig. 4i). This further demonstrates that language-related functional regions are associated with gene expression in module 11 and superficial L2-L3 glutamatergic types.

### Function-related laminar restrictions in non-neuronal cells

The atlas has comprehensive coverage of cortical non-neuronal cells, including 7 clusters of astrocytes (AST), 4 clusters of microglia (MICRO), and 5 clusters of oligodendrocyte progenitor cells (OPC). (Fig. 5a, b, and Extended Data Fig. 12). We observed the strong specific distribution of non-neuronal cell types in cortical regions and their similarity across regions by analyzing transcriptomic similarity analysis of non-neuronal cell types (Fig. 5c-d, and Extended Data Fig. 13). Besides, we found that OPCs, astrocytes, and microglia were most prominent in the distribution of the supramarginal gyrus, whereas oligodendrocyte was concentrated in the prefrontal cortex (Fig. 5c). Compared with non-human primates, humans exhibit a higher proportion of OPCs, suggesting that genes associated with the regulation of OPC maturation—specifically, *FOXP2*, *THEMIS*, and *PCDH15*—are more abundantly expressed in the human cortex^31^. Utilizing Stereo slides, we investigated the distribution of these genes and discovered that their highest overlapping regions are located within layer 6 and even in the WM (Fig. 5e). Moreover, our gene density analysis across multiple cortical regions revealed a likely higher prevalence of cells co-expressing *FOXP2* and *THEMIS* in the SPL (Fig. 5f).

**Fig. 5.**
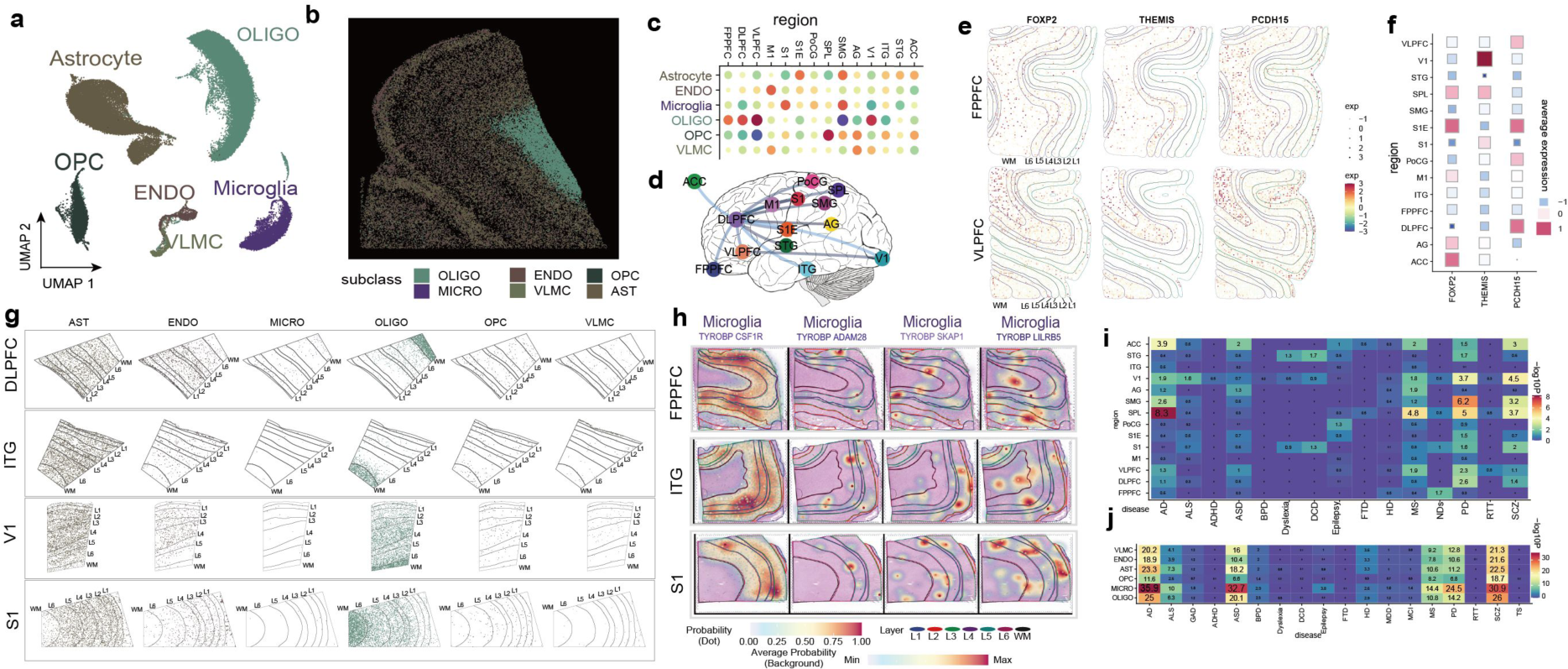
Transcriptomic taxonomy of non-neuron and spatial arrangement in human cerebral cortex. **a** UMAP visualization representing the clustering of different non-neuronal subclasses, including Astrocytes (AST), Oligodendrocytes (OLIGO), Endothelial cells (ENDO), Microglia (MICRO), Oligodendrocyte precursor cells (OPC), and Vascular mural cells (VLMC). **b** The cell type annotation for 37.5 μm bins of Stereo-seq section of human DLPFC by snRNA-seq at subclass level with only expression pattern of non-neuronal cells. **c, d** Regional transcriptomic similarities between cortical regions. Heatmap of the confusion matrix showing the separability of non-neuronal subclasses from different cortical regions (**c**). Constellation plots of regional transcriptomic similarities between cortical regions. The link between the two regions delineated a shared cell proportion in KNN discrimination with the region as a label and transcriptome as a feature (**d**). Only the link with shared fraction over 5% was shown. **e** Stereo spatial slides of prefrontal lobe showing expression of three genes related to brain evolution. **f** Heatmap showing the expression of three genes related to brain evolution in each cortical region. **g** Laminar distribution of non-neuronal subclasses at the Stereo spatial cortical regions. Dot plots show the Stereo spatial transcriptome results of the non-neuronal cells divided into 6 different subclasses. The color and alpha of spots denote the position of each cell. **h** Decomposition of cell type mixtures in spatial transcriptome and divided into 4 Microglia cell types according to the function and expression patterns. Color and alpha denote the average cell-type probabilities around spots. **i, j** Heatmap showing the enrichment of neural disease gene set in different non-neuronal transcriptomes across multiple cortical regions (**i**) and non-neuronal subclasses in human traits (**j**). AD, Alzheimer’s disease; ALS, Amyotrophic lateral sclerosis; GAD, Generalized anxiety disorder; PPA, Primary progressive aphasia; ADHD, Attention deficit hyperactivity disorder; ASD, Autism spectrum disorder; BPD, Bipolar disorder; NPD, Neuronal plasticity disorder; Dyslexia; DCD, Dyspraxia; Epilepsy; FTD, Frontotemporal lobar degeneration; HD, Huntington’s disease; MDD, Major depression disorders; MCI, Mild cognitive impairment; MS, Multiple sclerosis; NDs, Neurodevelopmental disability; PD, Parkinson’s disease; PTSD, Post-traumatic stress disorder; RTT, Rett syndrome; BCECTS, Rolandic epilepsy with speech impairment; SCZ, Schizophrenia; TS, Tourette syndrome.

Most non-neuronal clusters differ in layer distribution, which relates to their specific expression patterns and functions restricted to specific sites, such as guiding neuronal migration or stabilizing synaptic connections (Fig. 5g). From the more intuitive Stereo-seq of multiple cortical regions, the distribution of oligodendrocytes (OLIGO) is more inclined to WM, while OPC is relatively scattered at other levels, even though their expression patterns are very similar (Fig. 5b, g). In view of our observations of large-scale multi-cortical regions, the OLIGO is stable in WM which is similar in MERFISH^32^. MICRO, in contrast, were dispersed from layer 1 to layer 6, and Vascular mural cells (VLMC) clusters occupied the supragranular layers (Fig. 5g).

We propose the transcription continuum model as a more effective way to classify microglia, considering their diverse functions and roles in brain transcript properties^33,34^. Under this model, MICRO fall into four distinct cell types based on their expression profiles: TYROBP CSF1R signifies neutral microglia; TYROBP ADAM28 characterizes proliferative zone-associated microglia; TYROBP SKAP1 identifies axon bundle-associated microglia; and TYROBP LILRB5 represents white matter-associated microglia (Fig. 5h). Classification of MICRO using the transcriptional continuum model takes into account their different functional and regional roles in the molecular properties of brain transcripts, but does not elucidate the role that MICRO play in the cortical immune microenvironment^35,36^.

Furthermore, we examined the susceptibility of non-neuronal subclasses in various cortical regions to diseases. The supramarginal gyrus showed the highest susceptibility to Parkinson’s disease, while microglia exhibited strong associations with multiple neuropsychiatric conditions (Autism spectrum disorder, Parkinson’s disease, and Schizophrenia) (Fig. 5i-j). Additionally, VLMC, Endothelial cells (ENDO), and MICRO in SPL were identified as particularly sensitive to Alzheimer’s disease (Extended Data Fig. 14).

### Development-related expression patterns and spatial arrangements of GABAergic neurons

GABAergic neurons are primarily involved in local communication within a specific brain region. They form connections with other neurons within the same brain region and play a critical role in the integration and processing of information within that region^37^. GABAergic neurons in the taxonomy were split into two groups: caudal ganglionic eminence (CGE)-derived and medial ganglionic eminence (MGE)-derived (Extended Data Fig. 15-16). In detail, we identified 4 subclasses of the CGE branch: LAMP5, NDNF, PAX6, and VIP (Fig. 6a). On the contrary, MGE-derived neurons have only two subclasses, parvalbumin (PVALB) and somatostatin (SST) (Fig. 6a), with developmental fates derived from ventral and dorsal MGEs, respectively^38,39^. In the adult human cortex, the CGE-derived and MGE-derived neurons accounted for 45% and 55% of the total (Supplementary Table 2), consistent with initial neurodevelopmental origins^4,13,40,41^, indicating a relatively stable composition of GABAergic neurons during the whole human life. Except for the specific distribution of LAMP5 in the ACC and the higher distribution of NDNF in the temporal lobe STG, most other GABAergic neurons were present in all 14 cortical regions, lacking significant encephalic regional specificity (Fig. 6b-c).

**Fig. 6.**
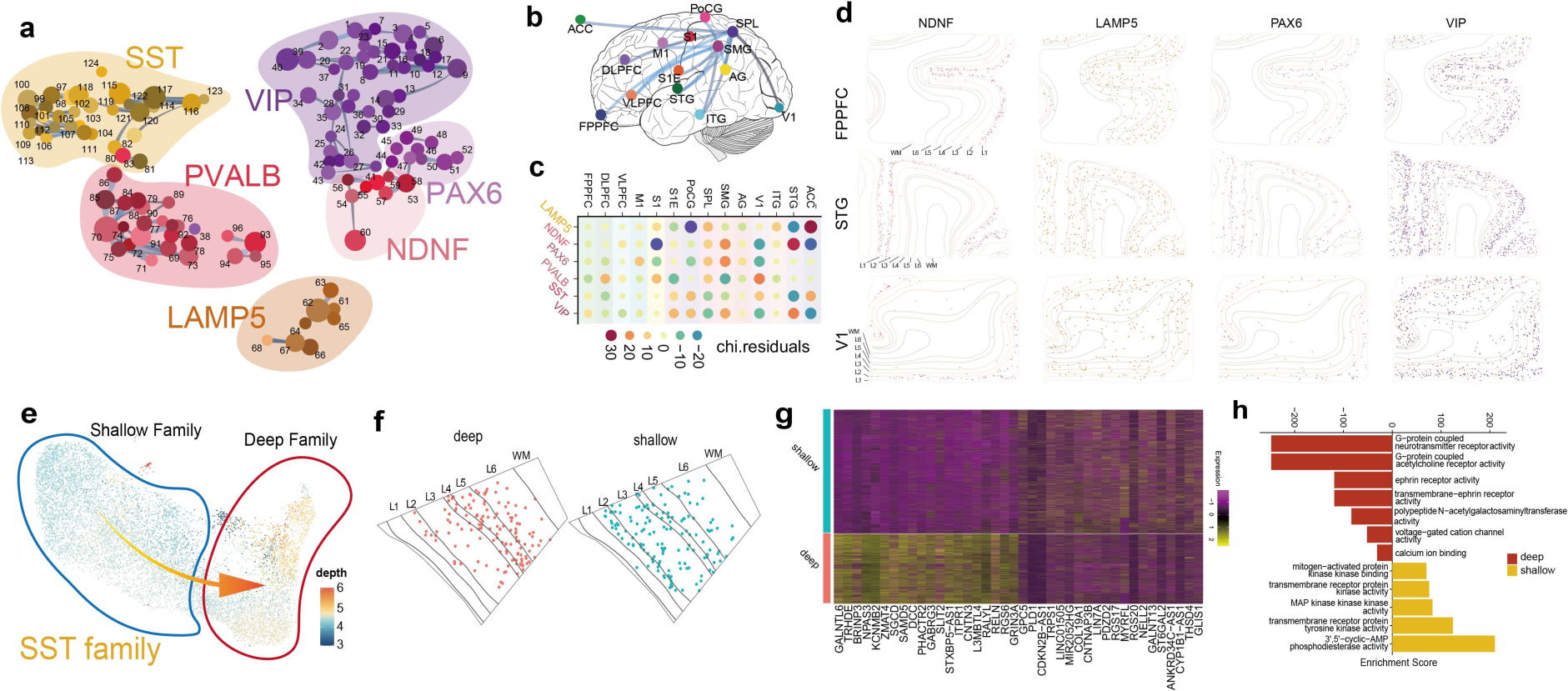
Characterization and spatial distribution of GABAergic subclasses in human cortical regions. **a** Constellation plots of the global relatedness of GABAergic subclasses, characterizing the relatedness among all clusters in different subclasses. Each subclass is represented by its specific color region, placed at the cluster centroid in UMAP. Each dot denotes a cluster within its subclass. **b, c** Regional transcriptomic similarities between cortical regions. Constellation plot of regional transcriptomic similarities between cortical regions. Separability of GABAergic neurons inferred using confusion matrices, displayed using associated stereo-similarity prediction plots (**b**). Only the link with shared fraction over 5% was shown. Heatmap of the confusion matrix showing the separability of GABAergic neurons from different cortical regions (**c**). **d** Distribution of typical CGE-derived subclasses at the Stereo spatial cortical regions. Dot plots show the Stereo spatial transcriptome results of the CGE-derived cells divided into 4 different subclasses. The color and alpha of spots denote the position of each cell. **e** UMAP plot distinguishing between the shallow and deep families within the GABAergic SST neuron, denoted by blue and orange, respectively. **f** Zoom-in Stereo slides view for deep and shallow SST family in cortical regions, illustrating difference distribution of MGE-derived deep and shallow SST family, with color of spots denote the position of each cell. **g** Heatmap of top 25 differential genes expression in deep and shallow SST family. **h** The enrichment of biological process in deep and shallow SST family.

In contrast to earlier genetic destiny^42^, taxonomic NDNF is present at the superficial layer. We found that most CGE-derived clusters favored a supragranular arrangement, whereas MGE-derived clusters had no apparent laminar restriction (Fig. 6d, and Extended Data Fig. 15-16). Compared with GABAergic neurons in the mouse cortex, the CGE-derived neurons in the human cortex migrate more shallowly within the tissue and contribute more cells that not only localize exclusively around layer 5^43^.

Through Stereo slides, it was found that MGE-derived SST neurons have upper and lower layering characteristics (Fig. 6e), and MERFISH also confirmed this^32^. SST neurons exhibit a division into upper and lower regions, with the L3 region showing relative sparsity (Fig. 6f), a difference attributed to distinct migration paths. Shallow SST neurons might migrate through the MZ, while deep-layer neurons might migrate via the SVZ, resulting in clear layering. When comparing SST neurons derived from the migration paths of the two groups of mice, MZ group exhibited upregulation in the expression levels of 24 genes compared to the SVZ group^44^. Additionally, we conducted a genetic differential analysis of superficial and deep MGE-derived SST neurons and found that 25 genes exhibited upregulation in deep SST neurons compared with superficial SST neurons (Fig. 6g). Furthermore, deep SST neurons are associated with G-protein coupled neurotransmitter receptor activity within their pathway, while superficial SST neurons are linked to cyclic adenosine monophosphate (cAMP) phosphodiesterase activity (Fig. 6h). We executed KEGG and GO term analysis to evaluate the substantial differences within GABAergic subclasses (Extended Data Fig. 17-18). Moreover, by overlaying neural risk gene sets across various cortical regions and subclasses of GABAergic neurons, we discovered a strong association between psychiatric disorders and the susceptibility of GABAergic neurons (Extended Data Fig. 19).

### Spatial variation of RORB glutamatergic neurons around layer 4

The cytoarchitectural atlas of the adult human cortex empowers us to locate layers or search for finer cortical structures. Previous studies suggested that RORB cells are selectively vulnerable neurons, notably L4-like neurons^45^, and traverse the supragranular and subgranular layers^46^. Although the previous study had identified markers of L4 (RORB) and L4-like neurons in M1^14^, it remains largely unknown how RORB cell types distribute across multiple human cortical regions. We proceeded to analyze the distribution density of all subclasses within each cortex (Fig. 7a). Our findings revealed that, within the gradient-laminar distribution of glutamatergic subclasses, L3-L4 IT and L4-L5 IT RORB glutamatergic neurons dominate in V1 (Fig. 7b, c). This prevalence appears to be linked to the visual cortex, and it is associated with the thicker layer 4 situated in the middle (Fig. 7d, e). Simultaneously, we also investigated the hierarchical structure of V1 in stereo slides and confirmed that the increased thickness of the granular layer accounts for the greater presence of RORB glutamatergic neurons (Fig. 7f, g).

**Fig. 7.**
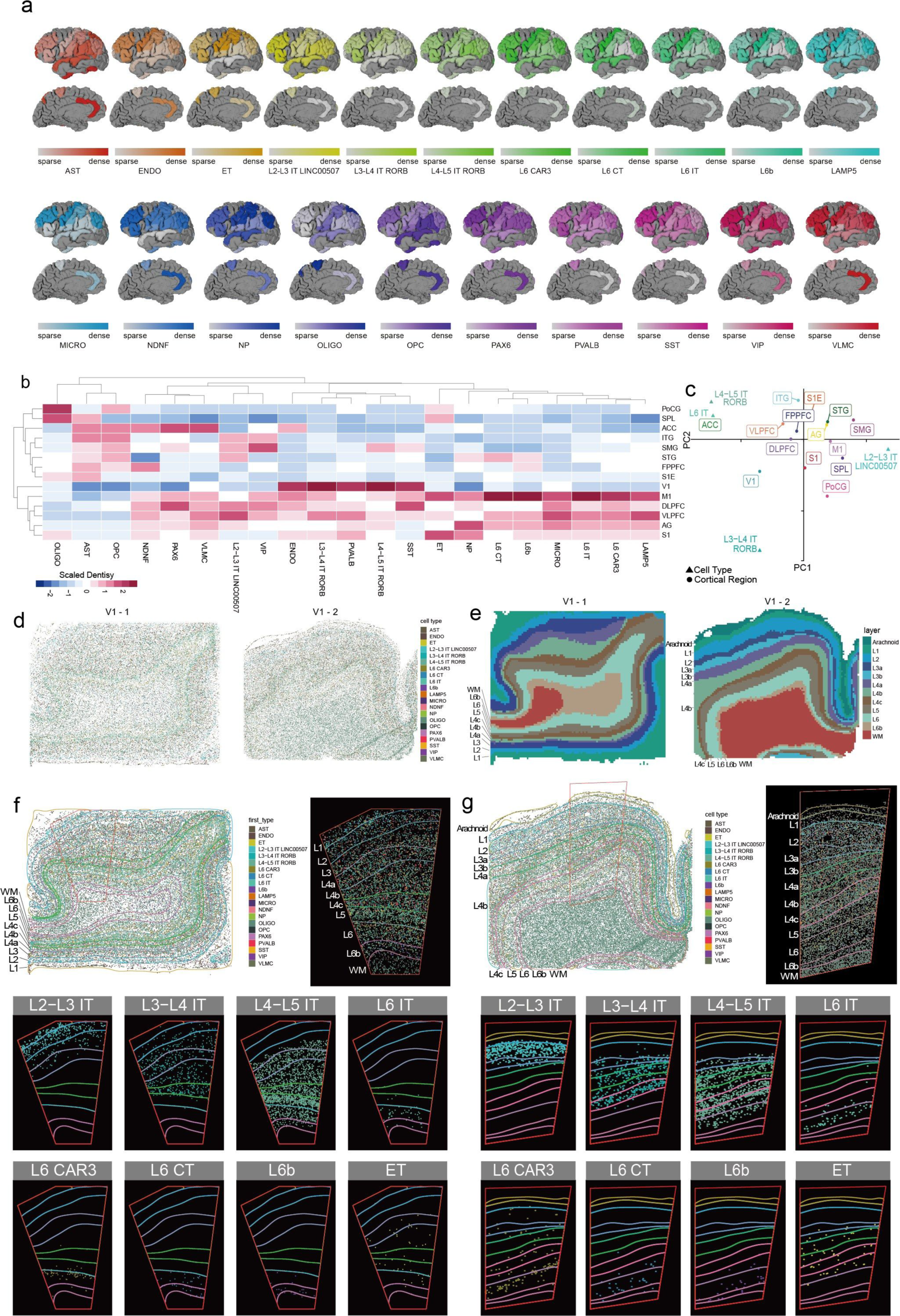
Spatial variation of RORB glutamatergic neurons in visual cortex. **a** The density distribution of each subclass in the cortical regions. The density statistics are shown below. The color follows the color assigned to each subclass in the taxonomy. **b** Heatmap showing the density distribution of each subclass in cortical areas. **c** The biplot showed the correlation between cortical region and glutamatergic subclasses by redundancy analysis (RDA). The angle reflects the correlation between cortical region and glutamatergic subclasses, and the length indicates the magnitude of each explicative contribution. **d** Cell-type annotation, subclass neuronal cell types was determined by clustering and RNA expression **e** Refined laminar annotation identification of the layer 4 in visual cortex, DeepST AI-assisted semi-supervised laminar segmentation based on nucleic acid and cell type annotation. **f, g** The spatially resolved single-cell transcriptome of the adult human telencephalon V1 as determined by Stereo-seq analyses (top left). Selected an area with obvious layered structure (top right). Scatterplot of two Stereo-seq example (V1) showing gradient-laminar distributed subclasses (down). Cells are colored by their subclass ID (same color as in a). The slide size is 1×1cm.

### De novo transcriptomic identification of the subplate area in human cerebral cortex

In addition to layer 4, we leveraged the atlas to search for finer structures in human cerebral cortex, like layer 6b (Fig. 8a). This structure was transcriptionally defined as layer 6b in the human cerebral cortex, and its neurons differ from those in layer 6 and express markers of cortical subplate neurons such as *CTGF (CCN2), SEMA3E, and MGST1, etc* (Fig. 8b, and Supplementary Table 4)^47,48^. These findings demonstrate that layer 6b is not a transient structure and is stabilized in the posterior segment of layer 6 versus the anterior end of the WM^49,50^, suggesting that cortical subplate neurons may be a permanent subpopulation. Cortical subplate neurons have only L6b in layer 6b to regulate the activity of deep cortical circuits^51^, while L6CT is a transitional cell type that exists between layer 6 and layer 6b (Fig. 8c).

**Fig. 8.**
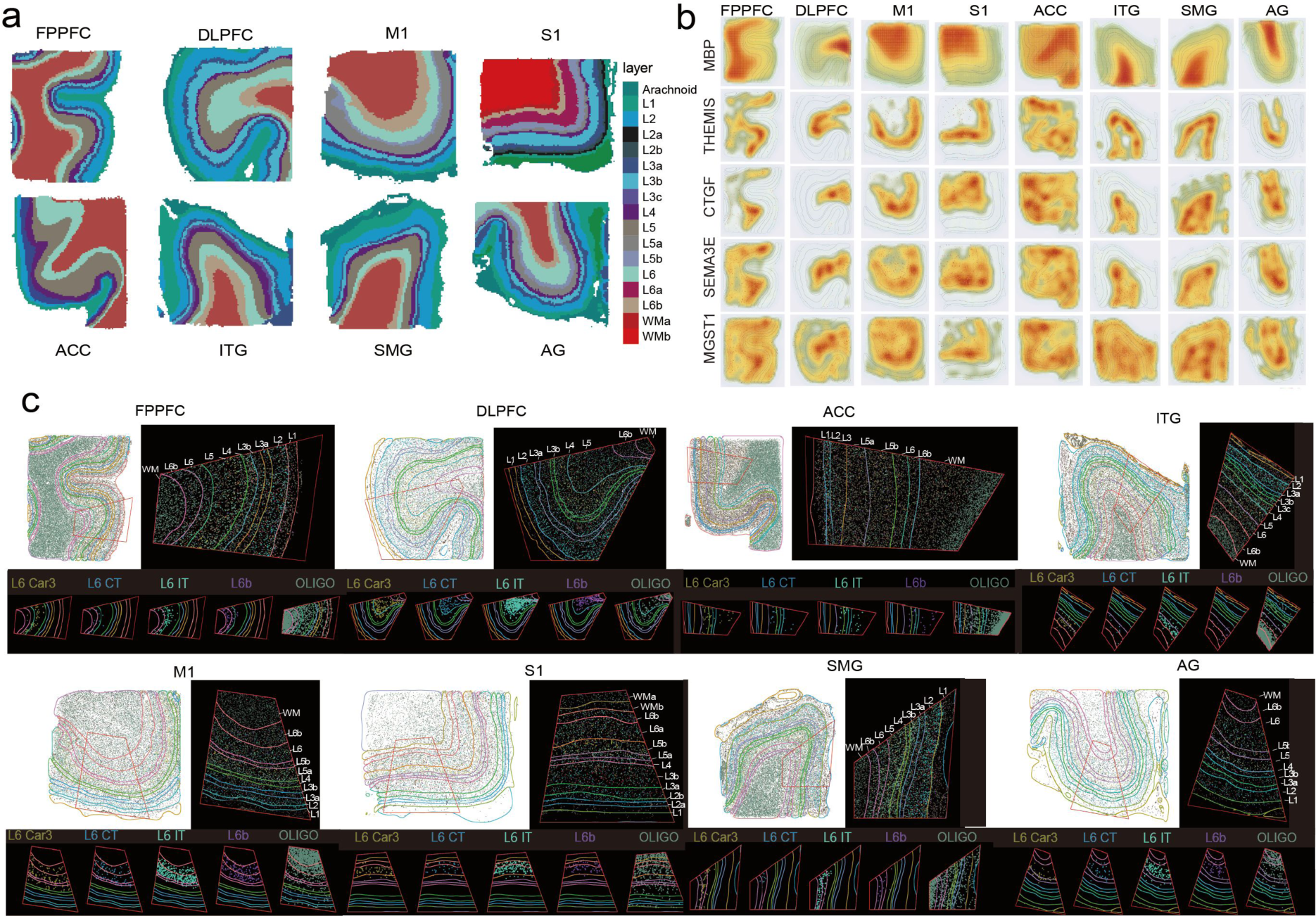
Transcriptomic identification of the layer 6b in human cerebral cortex. **a** Refined laminar annotation identification of 8 non-visual cortices, DeepST AI-assisted semi-supervised laminar segmentation based on nucleic acid and cell type annotation. **b** The expression of l6b markers in different regions. The expression is plotted by weighted expression density calculated by same genes expressed from adjacent spatial spots. **c** The spatially resolved single-cell transcriptome of the 8 cortical regions as determined by Stereo-seq analyses (top left). Selected an area with obvious layered structure (top right). Scatterplot of 8 Stereo-seq examples showing gradient-laminar distributed subclasses around layer 6 and white matter (down). Cells are colored by their subclass ID (same color as in Fig.7a). The slide size is 1×1cm.

## Discussion

We profiled the transcriptomes of 307,738 nuclei from 42 fresh samples, 46,948 VISIUM spots for 12 10x slides, and 1,355,582 cells for Stereo 32 slides of 8 donors across the entire human cerebral cortex, conducting a comprehensive census of cellular diversity and spatial arrangements, which facilitated the delineation of the cellular layer distributions of cell types and provided much more spatial and transcriptomic information than traditional approaches such as RNAscope^4,14,52^.

Our analysis shows that neural cells in the human cortex exhibit high diversity in cell types and heterogeneity in spatial arrangements. There are favorable mappings among different cellular attributions such as transcriptomically taxonomic classification, molecular features, laminar distribution, and developmental fate. We showed that differences in expression across different types of glutamatergic neurons may help explain the range of projection patterns (Fig. 3). Glutamatergic neurons appeared to be more layer-specific with unimodal distribution, like the RORB neuron type spanning the supra- and subgranular layers and the LINC00507 neuronal type prevalent in the superficial layer. Besides, four types of CGE-derived interneurons were found that tend to arrange in the superficial layer and are uniformly distributed throughout the cortex (Fig. 6c, d). Moreover, our atlas revealed that the localization of layer 4 and layer 6b exposes particular marker genes and cellular architecture.

Our cytoarchitectural atlas covers the entire human cerebral cortex, and samples with high quality and completeness improved the reliability and robustness of this work. Yet a larger cohort of samples with more cells and genes by more advanced techniques can promote more appropriate and accurate output, which helps find out rarer cell types and bring more novel insights into potential heterogeneities like genders and ages. Moreover, we took the robust clustering but conservative in spatial domain identification, which may ignore variability within clusters. Futural work should flesh out these differences, as well as link transcriptomic characteristics with morphology and electrophysiology^53^.

Overall, our study demonstrates the potential power of combining snRNA-seq and spatial transcriptome sequencing for exploring the cellular basis of sensory perception, cognition, and behavior. Research along these lines may not only help elucidate the mysteries of human cognition but also accelerate our understanding of neurological disorders^54^.

## Supporting information

Supplementary Table 1 Sample information and mRNA alignment statistics,

Supplementary Table 2 Laminar markers

Supplementary Table 3 Regional expression difference among subclass, pathway enrichment,and disease susceptibility

Supplementary Table 4 Data of reclustering on spots from WM and layer 6

## Acknowledgments

We thank NovelBio Bio-Pharm Technology Co., Ltd. (Shanghai, China) for the support of single-cell nucleus sequencing and spatial transcriptome sequencing (NovelBrain Cloud Analysis Platform, www.novelbrain.com).

## Author Contributions

Conceptualization: Q.J., and J.X; snRNA-seq and spatial transcriptomic data generation: S.W., M.L., P.W., X.J., and Q.J.; Data analysis: S.W., M.L., P.W., X.J., C.X., X.L., Z.X., H.L., and Q.J.; Data interpretation: S.W., M.L., P.W., and Q.J.; Write manuscript: S.W., M.L., and Q.J.

## Data and Materials Availability

All sequencing data could be downloaded with NGDC accession number PRJCA009779 under permission from the corresponding author. All codes are available on https://github.com/JiangBioLab/Human-Brain-Cytoarchitectural-Atlas. The raw data derived from the immunohistochemical staining process is accessible on figshare, under DOI: 10.6084/m9.figshare.24279433.

## Methods

### Human brain tissue acquisition

This study was approved by the Ethical Committee of the Harbin Institute of Technology (Approval No. HIT-2021004), and the Ethics Committee of Foshan Maternity & Child Healthcare Hospital (Approval No. FSFY-MEC-2021-129). Tissues were collected from individuals who died suddenly and had no known history of brain injury or neurological disease. All experiments were performed by following the guidelines of the International Review Board and Institutional Ethics Committee. Brain samples were collected after donor family members signed informed consent in compliance with sample use and storage guidelines. In addition, tissues were used with the informed consent of immediate family members. Coronal sections were used only if they were confirmed not to show neurological or neuropsychiatric disease, based on an assessment by a forensic expert in the Department of Forensic Medicine of Guangdong Medical University. All five coronal sections in the study were collected within 48 hours after death and showed an RNA integrity number (RIN) of at least 6.2. Besides, samples for spatial transcriptomic sequencing own RIN over 7 (Supplementary Table 1). The eight donors were 50 ± 13 years old at death.

### Brain tissue processing and staining

Coronal brain sections were suspended in pre-chilled to −80°C, oxygenated artificial cerebrospinal fluid [0.5 mM calcium chloride (anhydrous), 25 mM D-glucose, 20 mM HEPES, 10 mM magnesium sulfate, 1.2 mM sodium phosphate monobasic monohydrate, 92 mM *N*-methyl-D-glucamine chloride (NMDG-Cl), 2.5 mM potassium chloride, 30 mM sodium bicarbonate, 5 mM sodium D-ascorbate, 3 mM sodium pyruvate, and 2 mM thiourea]. The sections in artificial cerebrospinal fluid were quick-frozen on dry ice and transported to a pathology dissection table. The following cortical regions were identified in all five brain sections: frontopolar prefrontal cortex (FPPFC), dorsolateral prefrontal cortex (DLPFC), primary motor cortex (M1), primary somatosensory cortex (S1), primary visual cortex (V1), and inferior temporal gyrus (ITG). These regions were separately extracted from the five sections, cut into smaller pieces, immersed in OCT fixative (catalog no. 4583, Tissue-Tek® Sakura, Torrance, CA), quick-frozen on dry ice, and stored at −80 °C until cryosectioning.

The OCT-embedded tissue was cryosectioned (Leica CM1950, Heidelberger, Germany) to a thickness of 10 μm and placed pre-chilled to −20°C, VISIUM Tissue Optimization Slides (catalog no. 3000394, 10x Genomics) and VISIUM Spatial Gene Expression Slides (catalog no. 2000233, 10x Genomics). The backs of the slides were warmed up at 10°C for 5 seconds to allow the tissue to adhere. The sections were then fixed with pre-chilled to 0°C methanol and stained according to the VISIUM Spatial Gene Expression User Guide (version CG000239 Rev A, 10x Genomics) or VISIUM Spatial Tissue Optimization User Guide (CG000238 Rev A, 10x Genomics).

Sections were stained with hematoxylin-eosin as follows. Sections were incubated for 1 min at 37 °C, fixed in methanol at −20 °C for 30 min, incubated for 7 min in hematoxylin, for 2 min in Bluing Buffer (catalog no. C0105S, H&E staining reagent test kit, Beyotime, Shanghai, China), and 1 min in eosin. Between each staining step, slides were washed with DNase- and RNase-free water (catalog no. 10977023, Invitrogen™, MA, USA). Stained sections were imaged under a microscope (ECLIPSE Ti, Nikon, Tokyo, Japan).

Sections were stained with Nissl solution as follows. Sections were washed briefly in tap water to remove residual salts, immersed in 2 changes in 100% ethanol (3 min each), defatted through 2-3 changes of 100% xylene (15 min each), and rehydrated in 100% ethanol for 10 min, washed in tap water, stained for 4-15 min with 0.1% cresyl violet Nissl staining solution (catalog no. G1432, Solarbio, Beijing, China), rinsed quickly in tap water, then washed in 70% ethanol.

When necessary, Nissl-stained sections were immersed for 2 min in differentiation solution (catalog no. G1432, Solarbio, Beijing, China), dehydrated through 2 changes of absolute ethanol (3 min each), and cleared through x2 of xylene, then allowed to dry in a fume hood. Cortical layers in the sections were visualized under a fluorescence dissecting microscope (Leica DM6000B, Heidelberger, Germany).

### Immunohistochemistry staining

Coronal sections adjacent to Stereo-seq chips were collected for IHC staining with NeuN (Proteintech 26975-1-AP, 1:10000), GAP43 (Abcam ab75810, 1: 3000), and MAP2 (Proteintech 17490-1-AP, 1:2500) antibodies. The 10-μm sections were mounted on gelatinized glass slides and baked to dry for 5 minutes at 37 °C. Afterward, the brain tissues mounted on slides were fixed with 4% paraformaldehyde in 0.1M phosphate buffer (PBS) for 10 minutes. Following three PBS washes, the sections underwent a 15-minute pre-incubation in 0.5% Triton X-100 in PBS, followed by a 1-hour incubation in a blocking solution comprising 10% normal goat serum and 0.1% Triton X-100 in PBS (0.1% PBST). Subsequently, the sections were incubated overnight at 4°C in 0.1% PBST containing the monoclonal antibodies NeuN, GAP43, and MAP2 polyclonal antibody.

After another three washes in PBS, the sections underwent a 30-minute incubation in 0.1% PBST containing 0.6% hydrogen peroxide to block endogenous peroxidase that might contribute to background staining. Following three additional PBS washes, the sections were incubated in PBS containing a biotinylated secondary antibody (1:200) for 2 hours. They were then washed three times in PBS and transferred to PBS containing the peroxidase conjugate from the Vectastain CBC kit (Vector Laboratories, Burlingame, CA). After a series of rinses in PBS, the sections were immersed in a solution of 0.05% 3-3’diaminobenzidine-4HCl (DAB, Sigma-Aldrich, St Louis, MO) and 0.05% hydrogen peroxide. Once the staining was complete, the sections underwent dehydration in increasing concentrations of ethanol, were cleared in xylene, and finally, coverslipped with DPX medium. Subsequently, the glass-mounted sections were scanned at 5× (0.88 μm/pixel) in a Zeiss scanner to generate images.

### Tissue processing for spatial transcriptomic sequencing

VISIUM Spatial Tissue Optimization Slides and Reagent Kit (10X Genomics) were used. Cortical sections on the slides were permeabilized using Permeabilization Enzyme for varying amounts of time. Fluorescent RT Master Mix was added, and tissues were removed using Tissue Removal Mix for 60 min at 56 °C. The sections were examined under a fluorescence microscope (ECLIPSE Ti, Nikon, Tokyo, Japan) to identify the best permeabilization time. Based on this optimal time, tissues were permeabilized for spatial transcriptomic sequencing using VISIUM Spatial Gene Expression Slides & Reagent Kits (10X Genomics).

After permeabilization, the slides were incubated in RT Master Mix for 45 min at 53 °C for reverse transcription, then with Second Strand Mix for 15 min at 65 °C to initiate second-strand synthesis. The resulting barcoded cDNA was transferred from the slides, purified, amplified, fragmented, A-tailed, ligated with adaptors, and subjected to index PCR. The final libraries were quantified using the Qubit High Sensitivity DNA assay (Thermo Fisher Scientific, Cleveland, OH, USA). Their size distribution was determined using a High-sensitivity DNA chip on a Bioanalyzer 2200 (Agilent, Santa Clara, CA, USA). All libraries were sequenced in 150-bp paired-end runs on an Illumina sequencer (San Diego, CA, USA).

Tissue sections for Stereo-seq were adhered to the Stereo-seq chip (generated by BGI, China) surface and incubated at 37℃ for 3 minutes. Then, the sections were fixed in methanol and incubated for 30 minutes at −20℃ before Stereo-seq library preparation. Where indicated, the same sections were stained with nucleic acid dye (Thermo Fisher, Q10212) and imaging was performed with a Motic Custom PA53 FS6 microscope prior to in situ capture at the channel of FITC.

### Single-nucleus RNA sequencing

Samples were prepared for snRNA-seq at NovelBio Bio-Pharm Technology (Shanghai, China). Tissue samples were surgically removed as the methods above and snap-frozen in liquid nitrogen, and then nuclei were isolated as described, with some modifications. Briefly, frozen tissue was homogenized in NLB buffer [250 mM sucrose, 10 mM Tris-HCl, 3 mM MgAc2, 0.1% Triton X-100 (Sigma-Aldrich, Saint-Louis, Missouri, USA), 0.1 mM EDTA, 0.2 U/μL RNase Inhibitor (Takara, Tokyo, Japan)]. Then nuclei were isolated on a sucrose gradient, adjusted to approximately 1000 nuclei/μL, and loaded into single channels to generate single-cell Gel Bead-In-Emulsions (GEMs) using the Chromium Controller Instrument (10X Genomics) and Chromium Single Cell 3’ Reagent Kit (version 3.1, 10X Genomics). After reverse transcription, GEMs were broken, and barcoded cDNA was purified, amplified, fragmented, A-tailed, ligated with adaptors, and subjected to index PCR. The final libraries were quantified using the Qubit High Sensitivity DNA assay (Thermo Fisher Scientific, Cleveland, OH, USA). The size distribution of the libraries was determined using a High Sensitivity DNA chip on a Bioanalyzer 2200 (Agilent, Santa Clara, CA, USA). All libraries were sequenced by Novaseq6000 (Illumina, San Diego, CA, USA) on a 150 bp paired-end run.

Coronal brain sections on slides were used for cell capture, barcoding, reverse transcription, cDNA amplification, and library construction using the Chromium Single Cell 3’ Reagent Kit (version 2, catalog no. 120237, 10x Genomics) according to the manufacturer’s instructions.

### Quantification and statistical analysis

Minimal sample sizes were not determined; instead, the sample used here was similar to that reported in previous studies. Data were not collected through randomization, nor were analysts blinded to the data conditions because all data represented a single set of conditions.

### Processing of single-cell sequencing data

Single-cell sequencing data were processed using the analysis pipelines in Cell Ranger (version 6.0.2, 10X Genomics). Reads were aligned against the human reference genome refdata-gex-GRCh38-2020-A, and feature-barcode matrices were generated using the *count* function in Cell Ranger. Expression levels were calculated as counts per million reads (CPM). In some analyses, CPM values were transformed by log_2_(CPM + 1). Genes were considered to be detected if CPM > 0. CPM values depended not only on absolute transcript number but also on gene length. Thus, CPM values for short, abundant transcripts could be similar to those for long, less abundant transcripts.

### Quality control

Low-quality cells in each sample were identified and removed using the *isOutlier* function in the program scatter in R, which identifies outliers based on median absolute deviation (MAD) from the median value of each metric across all cells. Cells were defined to be low-quality if 1) The cell library size (total UMI counts) is smaller than 3 MADs; 2) The number of detected genes is smaller than 3 MADs; 3) the proportion of mitochondrial gene counts is bigger than 3 MADs. Doublets were identified using DoubletFinder^55^, and the expected doublet rate was 0.075.

With the data from the remaining high-quality cells, we removed genes expressed in fewer than 10 cells to avoid unnecessary computational work in downstream analyses. Besides, 2 clusters were assigned to donor-specific due to a higher proportion of donors and were excluded from further analysis. These donor-specific clusters came from one donor in the dendrogram with very low reads and genes detected.

### Single-cell transcriptomic clusters

We adopted a mature iterative pipeline for clustering the same as with the previous studies^10^. Raw counts were log_2_-transformed (CPM + 1) values across nuclei, and results were clustered using the *consensus_cluster* function in the scrattch.hicat package in R and the parameters q1.th = 0.3, q.diff.th = 0.7, de.score.th = 80, and min.cells = 20. The iterative clustering was repeated 100 times on 80% of subsampled sets of cells, and the final clustering was based on a cell-cell co-clustering probability matrix. The average expression of marker genes at the cluster level was based on normalized gene expression, then constructed a dendrogram using the *build_dend* function in scrattch.hicat. Complete-linkage hierarchical clustering was performed using the *hclust* function in R based on default parameters. The resulting dendrogram branches were re-ordered such that excitatory clusters followed inhibitory clusters, and larger clusters appeared first while the tree structure was retained.

### Uniform manifold approximation and projection of single-cell data

We performed a principal component analysis based on the normalized gene expression matrix of 2000 highly variable genes. We used Harmony^56^ to perform batch correction across all samples, and then uniform manifold approximation and projection (UMAP) were performed with the *RunHarmony* function in Seurat^57^. Constellation plots depicting global relatedness among cell types were generated using the *get_knn_graph* and *plot_constellation* functions in scrattch.hicat^10^.

### Selection of markers of cell-type clusters

We used NS-Forest (version 2.1, RRID SCR_018348) to identify the minimum set of marker genes whose combined expression identified the cells of one type with maximum accuracy^58^ (https://github.com/JCVenterInstitute/NSForest). For each cluster, NS-Forest finds several combinations of a few binary markers. The top ten binary genes were identified for each cluster.

### Cluster annotation

Major cell types were first identified based on a common set of broad cell-type marker genes: expression of GAD1 or GAD2 identified GABAergic neurons; SLC17A7/SATB2, glutamatergic neurons; PDGFRA, oligodendrocyte progenitor cells; AQP4, astrocytes; PLP1/MOBP, oligodendrocytes; PDGFRB, vascular smooth muscle cells; FLT1, vascular endothelial cells; and APBB1IP, microglia.

We then defined subclasses and clusters based on the taxonomy dendrogram generated from data of multiple cortical regions. Following the convention of nomenclature^4^, Clusters were named according to the major cell type (EXC, INH, ASTRO, OLIGO, OPC, VLMC, and MICRO), the maximal expression subclass markers (*PAX6, LAMP5, VIP, SST, PVALB, LINC00507, RORB, THEMIS, FEZF2, TYROBP, FGFR3, PDGFRA, OPALIN* or *NOSTRIN*) and the top marker for each cluster.

### Regional similarity analysis

The analysis was done at the subclass level based on single-cell transcriptomics. We used randomForest on single-cell transcriptomic data for each category to build the classification model and obtain the appropriate confusion matrix, which can show how dissimilar cellular transcriptomics are from those from other regions. More comparable molecular features from different areas are represented by higher levels of confusion. The confusion matrix was then further clustered by *hclust* function to show the similarities in transcriptome profiles between various locations. Additionally, *FindAllmakers* in Seurat^57^ was used to examine the genes that differ in expression across regions.

### Functional analysis

The default mode network data of neurosynth in each region was used to access the correlation between functions of these regions of interest (ROI) and cell types. First, meta-analytic functional decoding analysis using NeuroSynth (www.neurosynth.org) database was conducted with NiMARE (https://github.com/neurostuff/NiMARE) to capture the most relevant cognitive functions. Specifically, the Neurosynth ROI association method and feature terms were downloaded from the 50 topic terms (v3; https://github.com/NeuroanatomyAndConnectivity/gradient_analysis/blob/master/gradient_data/neuro synth/v3-topics-50-keys.txt). Then, we combined these data with the corresponding cell type and subclass distribution data using the redundancy analysis function RDA from the vegan package. Hellinger transformation is used to standardize the data. The output data were scaled further using the function fortify from the ggplot2 package, with the scale parameter set to 2, which emphasizes the correlation between cell type/subclass and functions based on an angle rather than distance.

### Pathway enrichment and evaluation of heritability within cell-types

We used the Seurat function findMarker to determine the differentially expressed genes at the region, subclass, and region-subclass levels, using FDR as multiple corrections. The KEGG and GO terms enrichment based on the DEGs was accomplished with the help of a Clusterprolifier. We used the high-risk gene of various neurological diseases that was found in a recently established database for brain diseases BrainBase to evaluate the heritability at different levels. Following the process of filtering and unifying the diseases, our final goal was to target 27 disease sets with at least 10 genes for each type. The significance of the relationship between DEGs and disease-related gene sets was determined using the Fisher test.

### Data analysis and visualization of single-cell data

The single-cell transcriptomic data were analyzed or visualized using R (version 3.5.0 and higher, https://www.R-project.org), RStudio IDE (https://www.rstudio.com/), Seurat^57^, and scrattch suite (https://github.com/AllenInstitute/scrattch)^4,10^. All statistics tests without specifying were conducted by the Wilcox test.

### Validation with external cortical cell types

We compared the cortical cell types that we identified based on RNA sequencing with the cell types in transcriptomic atlases of M1 or M1 human primary motor cortex (https://portal.brain-map.org/atlases-and-data/rnaseq/human-m1-10x)^14^ and multiple human cortical regions (https://portal.brain-map.org/atlases-and-data/rnaseq/human-multiple-cortical-areas-smart-seq).

Median expression levels of cell-type marker genes were common to our results and these external datasets were used to map cells to a specific type. Mapping involved 100 iterations, during each of which 80% of genes were randomly selected, and the correlation between gene expression in the given cell with the median gene expression in each cell type in our dataset was computed. Based on the cell type identified during the 100 iterations, the probabilities that the given cell belonged to each of those types were calculated. The cell type that emerged as most probable was defined to be the type for the given cell.

### Spatial transcriptomics library generation

For VISIUM, libraries were prepared according to the VISIUM Spatial Gene Expression User Guide (<u>CG000239</u>). Libraries were loaded at 300 pM into a NovaSeq 6000 System (Illumina) and sequenced using a NovaSeq S4 Reagent Kit (catalog no. 20027466, Illumina) for 200 cycles at a depth of approximately 250-400 million read-pairs per sample. Sequencing was performed using the following read protocol: read 1, 28 cycles; i7 index read, 10 cycles; i5 index read, 10 cycles; read 2, 91 cycles.

For Stereo-seq, tissue sections placed on the chip were permeabilized with 0.1% pepsin (Sigma, P7000) in 0.01 M HCl buffer, incubated at 37°C for 5 minutes, and then washed with 0.1x SSC buffer (Thermo, AM9770) supplemented with 0.05 U/ml RNase inhibitor (NEB, M0314L) (NEB, M0314L). RNA released from the permeabilized tissue and captured by the DNB was reverse transcribed overnight at 42°C using SuperScript II (Invitrogen, 18064-014, 10 U/ml reverse transcriptase, 1 mM dNTPs, 1 M betaine solution PCR reagent, 7.5 mM MgCl2, 5 mM DTT, 2 U/ml RNase inhibitor, 2.5 mM Stereo-seq-TSO, and 1x First-Str After reverse transcription, tissue sections were washed twice with 0.1x SSC buffer and digested with Tissue Removal buffer (10 mM Tris-HCl, 25 mM EDTA, 100 mM NaCl, 0.5% SDS) at 55°C for 10 minutes. cDNA-containing chips were then treated overnight at 55°C with Prepare cDNA Release Mix (enzyme, buffer). VAHTSTM DNA Clean Beads (0.8×) purified cDNA. The concentrations of the PCR products were measured using the Qubit TM dsDNA Assay Kit (Thermo, Q32854). After fragmenting 20 ng of DNA at 55°C for 10 minutes with in-house Tn5 transposase, the reactions were terminated by adding 0.02% SDS and mixing gently at 37°C for 5 minutes. This is how fragmented products were amplified: 25 ml of fragmentation product, 1x KAPA HiFi Hotstart Ready Mix, 0.3 mM Stereo-seq-Library-F primer, 0.3 mM Stereo-seq-Library-R primer, and nuclease-free H2O in a total volume of 100 ml. The reaction was then carried out as follows: 1 cycle of 95°C for 5 minutes, 13 cycles of 98°C for 20 seconds, 58°C for 20 seconds, and 72°C for 30 seconds, and 1 cycle of 72°C for 5 minutes. PCR products were purified with AMPure XP Beads (0.63 and 0.13), utilized for DNB generation, and then sequenced on an MGI DNBSEQ-Tx sequencer.

### Spatial transcriptomics data preprocessing

VISIUM Raw FASTQ files and images of brain sections stained with hematoxylin-eosin were preprocessed using Space Ranger software (version 1.0.0 with STAR v.2.5.1b 52 inside) to generate the expression matrix. The reference genome was the Cell Ranger hg38 reference genome “refdata-cellranger-GRCh38-3.0.0” (http://cf.10xgenomics.com/supp/cell-exp/refdata-cellranger-GRCh38-3.0.0.tar.gz). Data of spots were removed if Space Ranger detected there was no tissue from the image data and scatter packages were used to do QC for each spot.

An MGI DNBSEQ-Tx sequencer was used to generate Stereo-seq fastq files. CID and MID are present in read 1 (CID: 1-25 bp, MID: 26-35 bp), whereas read 2 contains the cDNA sequences. First, the CID sequences on the initial reads were mapped to the designed coordinates of the in situ captured chip obtained from the initial round of sequencing, allowing for 1 base mismatch to correct for sequencing and PCR errors. Reads with a MID containing N or more than 2 bases and a quality score below 10 were filtered out. Each read’s CID and MID were appended to the header of each read. Using STAR (Dobin et al., 2013), retained reads were aligned to the reference genome ensemble 93 and mapped reads with MAPQ > 10 were counted and annotated with their corresponding genes). UMI with the same CID and gene locus were collapsed, allowing for the correction of sequencing and PCR errors with a single mismatch. Finally, this information was used to generate an expression profile matrix containing CIDs. The entire process has been incorporated into the publicly accessible pipeline SAW, which can be found at https://github.com/BGIResearch/SAW.

### Spatial transcriptomics data processing

To integrate 11 datasets of VISIUM and get the integrated expression matric, we performed Seurat’s spatial workflow (https://satijalab.org/seurat/articles/spatial_vignette.html) on each slide. Raw expression matrices were introduced by the Load10X_Spatial function and placed into Seurat objects with accompanying metadata^57^. Mitochondrial genes were removed then and all Seurat objects with each slide were normalized by the *SCTransform* function separately. Then, we integrated their expression matrix by 1) finding their 5,000 high variance genes first by function *SelectIntegrationFeatures* function; 2) Find anchors spots in each dataset by the *FindIntegrationAnchors* function; 3) Integrated the matrix by *IntegrateData* function. All results of gene expression were based on these normalized data. For the boxplot, we calculated the average expression of each gene in a different layer and perform Wilcox statistic tests.

### Spatial domain identification and annotation

To identify the spatial domain robustly, we adopted a deep learning framework (https://github.com/spatial-Transcriptomics/DeepST), to integrate histological and transcriptomic data. First, we segmented images of brain tissue stained with hematoxylin-eosin according to spot coordinates to obtain a partial image, and tissue topography was first processed using a pre-trained deep learning network (CNN). Morphological similarity *MS_ij_* between adjacent spots was calculated using this matrix, and the weights of gene expression *GC_ij_* and spatial location *SW_ij_* were merged to re-assign a normalized expression value 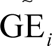 for each gene inside a spot S*_i_*The correlation was applied to calculate the weights of spatial gene expression using cosine distances GC*_ij_* between a spot S*_i_* and a spot S*j*:

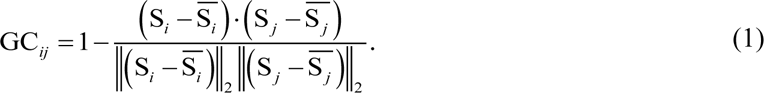

To represent the spot morphology, we used principal component analysis to extract the first 50 principal components as latent characteristics. The weights of morphological similarity MS*_ij_* between a spot S*_i_* and its adjacent spots S*j* were calculated using the cosine distance:

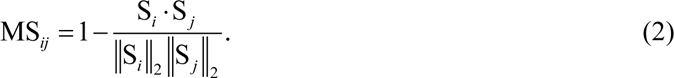

We used spatial coordinates to determine the Euclidean distance between each spot and the remainder, then ordered the distances between the top 3 (optional) adjacent spots to count the radius *γ* (mean add variance). For a given spot S*i*, a spot S*j* was a neighbor, so SW*_ij_* = 1 if and only if the Euclidean between two spots is less than the set value. Otherwise SW*_ij_* = 2.

We normalized the gene expression 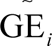 of each spot S*_i_* by incorporating gene expression correlation, spatial neighbors, and morphological similarity:

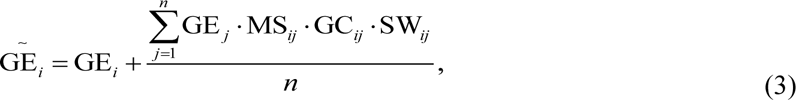

This also computed a graph adjacency matrix based on spatial coordinates by k-nearest neighbors (KNNs). The number of nearest neighbors was 15. Second, a denoising autoencoder was employed to perform nonlinear mapping on the integrated feature space, thereby generating a low-dimensional representation space that would reduce the risk of model overfitting. Simultaneously, a variational graph autoencoder was inserted into the framework to map spatial associations of spots, generating spatial embedding via integrated representation with the corresponding spatial adjacent spots.

The model was trained according to the following parameters: number of model training, 1000 epochs; the number of dimensions of the normalized gene expression matrix (*concat_pca_dim)*, 50; neural network type, graph convolutional neural network (*Conv_type=’GCNConv’*); and several integrated neighbors (*neighbour_k),* 2. Other parameters were default.

Finally, we obtained a 28-dimensional vector for each spot, for which we computed the Leiden clustering using Scanpy^59^. The clustering resolution varied adaptively with the number of spatial domains; it was 7 for human cortical regions.

Based on this primary result, we further corrected and annotated spatial domains manually in terms of their structural features, layer marker genes, Nissl-stained images, and cell type deconvolution.

### Cell-type proportion analysis

We deconvolute normalized spatial transcriptomic data with the transcriptomic spectrum of each cell type by RCTD^60^ and Tangram^61^ to achieve the robust results for VISIUM and Stereo-seq data. We selected the top 20 marker genes of pairwise 242 cell types and mapped all cell-type snRNA-seq profiles in spatial spots by totally 4637 genes. Default values were used for other parameter settings in all analyses. We calculated the probability of each cell type in each spot to calculate their subclass proportion in each pot to draw scatter pie plots. For the cell-type probability boxplot, we calculated the average probability of each cell type in different layers of each slide and performed Anova to measure global relatedness and the Wilcox test for specific pairwise comparison. For the laminar distribution and relative depth, we set 1∼7 as the relative depth value for layer 1 to white matter and calculated the standard deviation of probability based on these. For each cell type, we used kernel density estimation to smooth local probability to get weighted density by *kde2d.weighted* in package ggtern and visualized by *geom_raster* in ggplot2^62^.

### Spatial transcriptome UMAP

We used the previous algorithm to get the reduction of dimension by treating different samples as different batches and retaining default values for all other parameters. First, we normalized spatial gene expression by integrating spatial location information and tissue morphological information (see “Spatial domain identification” above) and then used principal component analysis to obtain a low-dimensional map of the normalized data, with a default of 50 dimensions. Second, the results were corrected for batch effects using Harmony^56^. The processed data were used to train the model, and these will generate a nonlinear, low-dimensional map of the integrated data. Finally, we applied the UMAP algorithm to this data to generate the 2-dimension projections.

### De novo clustering of spots from WM and layer 6

The previous algorithm for spatial domain identification offers robust but conservative results, which lose the sensitivity in tiny structures. To further explore the finer structure, we conducted transcriptomic clustering on spots annotated as WM or L6 from all 11 slides. These data were normalized by SCTtransform and further clustered by the default method in Seurat. The pipeline used the default parameters in the function *RunPCA* and *FindNeighbors* while the resolution is set to 0.1 in the function *FindClusters* to get a proper number of clusters. These allowed a finer clustering result derived from the previous annotations. We validated the new clusters by retrieving them into spatial slides and checking their specific gene expression and specific cell-type probability, and the result illustrates clusters with different reasonable biological contents and offers convincing evidence for their rationality.

### Data analysis and visualization of single-cell data

The single-cell transcriptomic data were analyzed or visualized using R (version 3.5.0 and higher, https://www.R-project.org), RStudio IDE (https://www.rstudio.com/), Seurat ^57^, and scrattch suite (https://github.com/AllenInstitute/scrattch) ^4,10^. All statistics tests without specifying were conducted by the Wilcox test.

## Extended Data Figures

**Extended Data Fig. 1.**
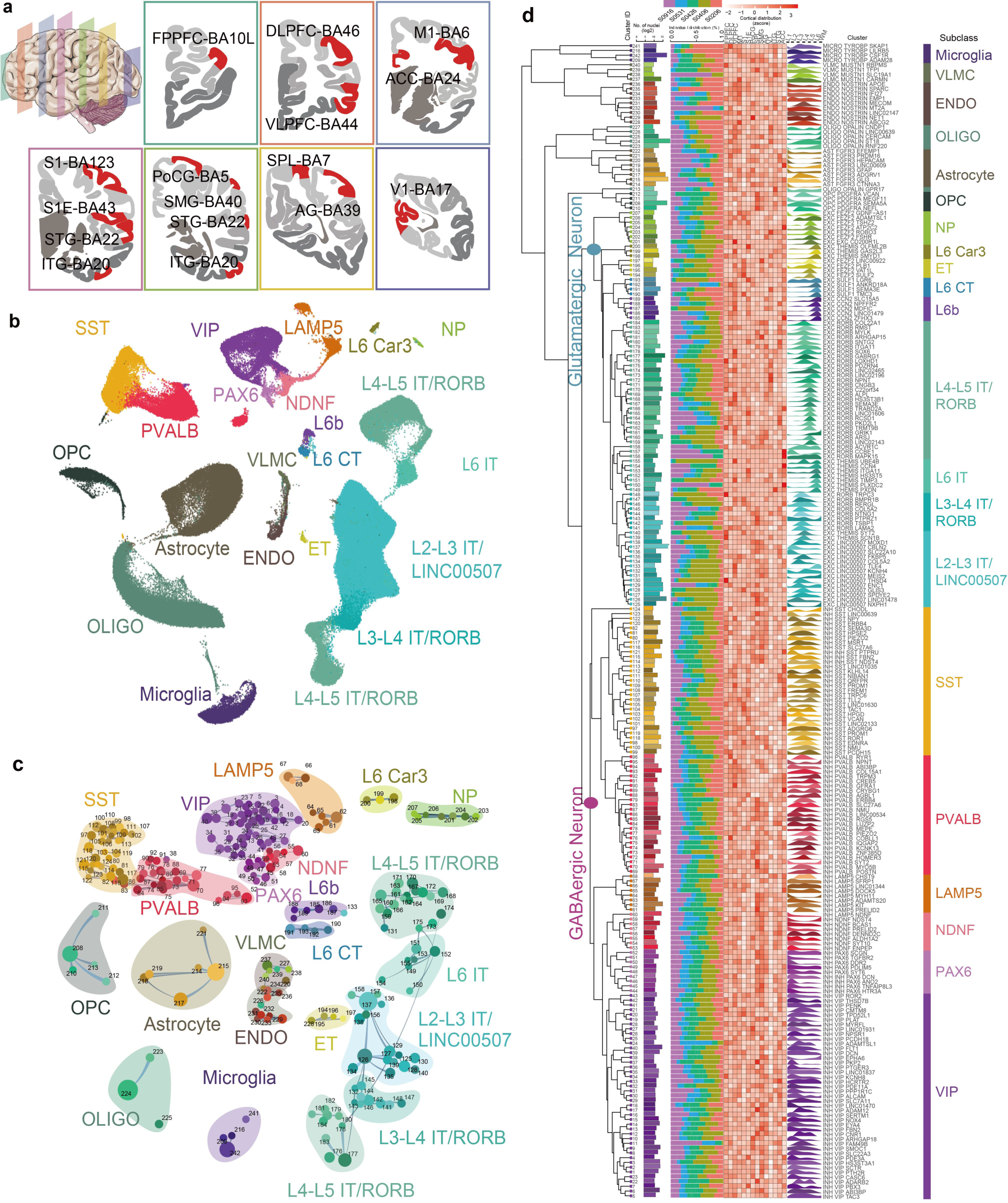
Transcriptomic diversity across the 14 adult human cortical regions. **a.** Overview of sampled cortical regions rendered in coronal sections. The samples cover the following 14 cortical regions. Colors denoted different coronal cortical sections, and the red color shows the part for dissection. **b.** UMAP representation of all subclasses in snRNA sequencing. Subclasses are labeled with different colors uniformly in all figures. **c.** Constellation plots of the global relatedness of all cell types, characterizing the relatedness among 242 clusters in different subclasses. Each subclass is represented by its specific color region, placed at the cluster centroid in UMAP. Each dot denotes a cluster within its subclass. **d.** The transcriptomic taxonomy of 242 clusters organized in a dendrogram. Most subclasses of cell types present clear preferences in spatial distribution and expression patterns. Bar plots showed the cell number of each cell type at log10 level (left). Percentage plot and heatmap represent fractions of cells profiled according to the donor and cortical region (middle). The ridge plot outlined the spatial arrangement of all 242 neural types across regions by convolution of the spatial transcriptome with snRNA-seq data (right).

**Extended Data Fig. 2.**
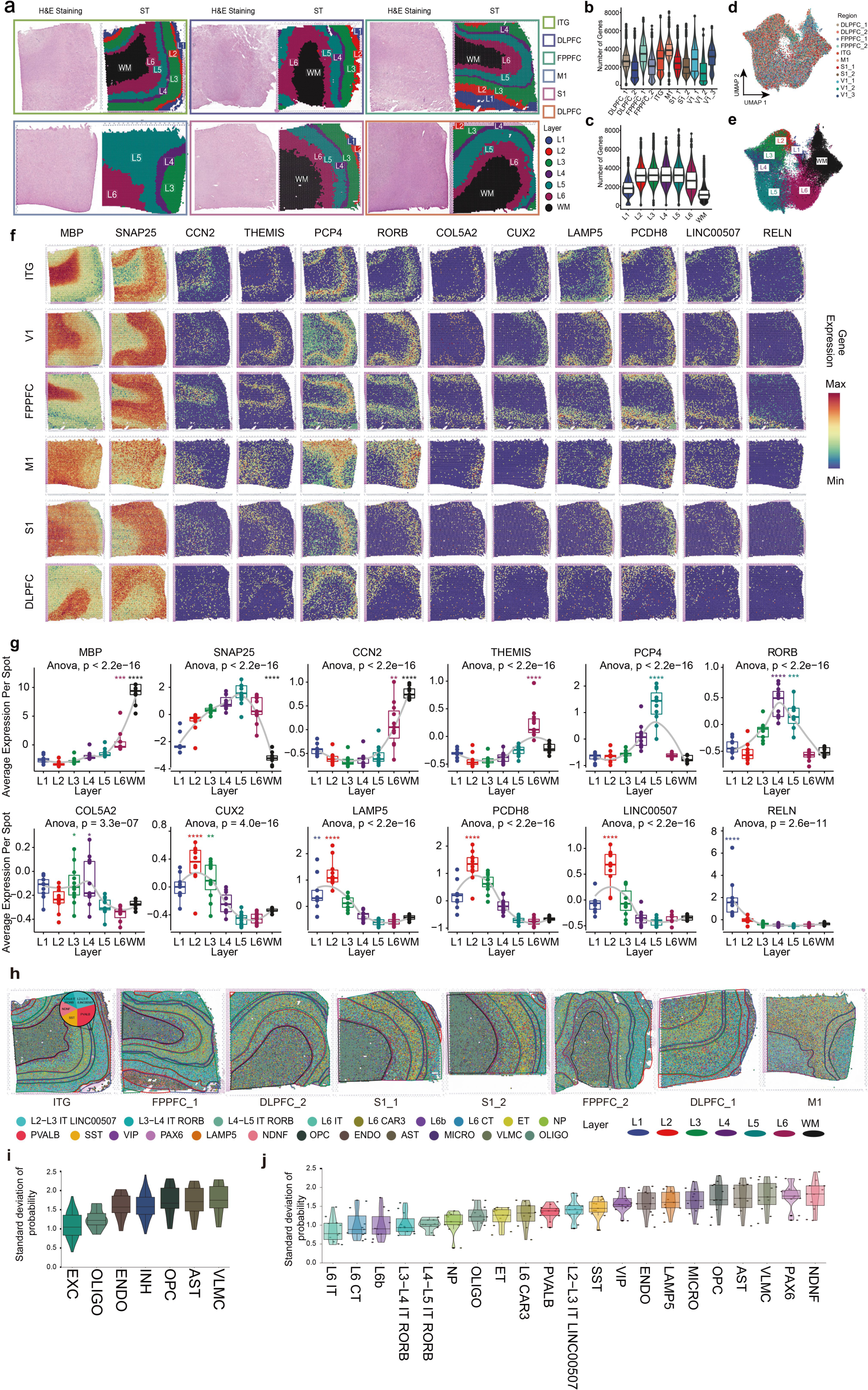
VISIUM slides of 5 regions and distribution of layer enrichment of taxonomic subclasses. **a.** Spatial transcriptome sequencing was performed on 6 cortical regions. Left, H&E staining; Right, Identification of laminar spatial domains (Methods). The slide size is 6.5mm × 6.5mm. Each slide has completed transcriptomic segmentation mostly with L1 to WM. We captured a total of 46,948 spots from 11 slides. **b.** The number of genes detected in each sample. Most median gene numbers per spot of samples are over 2,000 and the global median gene number of 2,605 per spot. **c.** The number of genes detected in each layer across 11 slides. These differences between L1/WM versus other layers reflect the high level of cellular density and activity in the middle cortical layers. **d, e.** UMAP representation of spots colored by the sample and layer annotation in integrated data of 11 slides. **f.** Spotplots of normalized expression for sample ITG, V1, M1, FPPFC, S1, and DLPFC for genes MBP (marker for WM), SNAP25 (marker for Neuron), CCN2 (marker for layer 6b), THEMIS (marker for layer 6), PCP4 (marker for around layer 5), RORB (marker for around layer 4), COL5A2 (marker for layer 4 and layer 3), CUX2 (marker for layer 3 and layer 2), LAMP5 (marker for layer 2), PCDH8 (marker for layer 2), LINC00507 (marker for layer 2) and RELN (marker for layer 1). **g.** Boxplots of median gene expression per layer across 11 slides. MBP (WM > rest, P = 1.57e−06), SNAP25 (layer 1 to layer 6 > rest, P = 2.61e-06), CCN2 (WM > rest, P = 3.08e−06 layer 6 > rest, P = 1.49e−03), THEMIS (layer 6 > rest, P = 1.58e−07), PCP4 (layer 5 > rest, P = 3.35e−07), RORB (layer 5 > rest, P = 3.81e−04; layer 4 > rest, P = 3.89e−04), COL5A2 (layer 4 > rest, P = 2.32e−02; layer 3 > rest, P = 1.66e−02), CUX2 (layer 3 > rest, P = 3.78e−03; layer 2 > rest, P = 5.98e−05), LAMP5 (layer 2 > rest, P = 1.76e−06; layer 1 > rest, P = 4.46e−03), PCDH8 (layer 2 > rest, P = 3.34e−06), LINC00507 (layer 2 > rest, P = 9.93e−07) and RELN (layer 1 > rest, P = 1.57e−06), related to G. For each layer, we calculated their pseudo-bulked gene expression for each layer of each sample. Significance was calculated by Wilcox test, p<0.05: *; p<0.01: **; p<0.001: ***; p<0.0001: ****. **h.** Spatial cellular compositional analysis in each spot shows all the taxonomic subclasses’ positional predictions. Cells are colored by their subclass identities (same colors as in Extended Data Fig.1). The slide size is 6.5mm × 6.5mm. With pie charts, the magnification of an ST-spot shows its subclass composition. **i.** Boxplot of deviation of major cell types probabilities show spatial laminar concentration. Smaller value illustrates higher density and compactness in spatial distribution while bigger value means wider and higher comprehensive distribution. **j.** Boxplot of deviation of subclass cell types probabilities show spatial laminar concentration.

**Extended Data Fig. 3.**
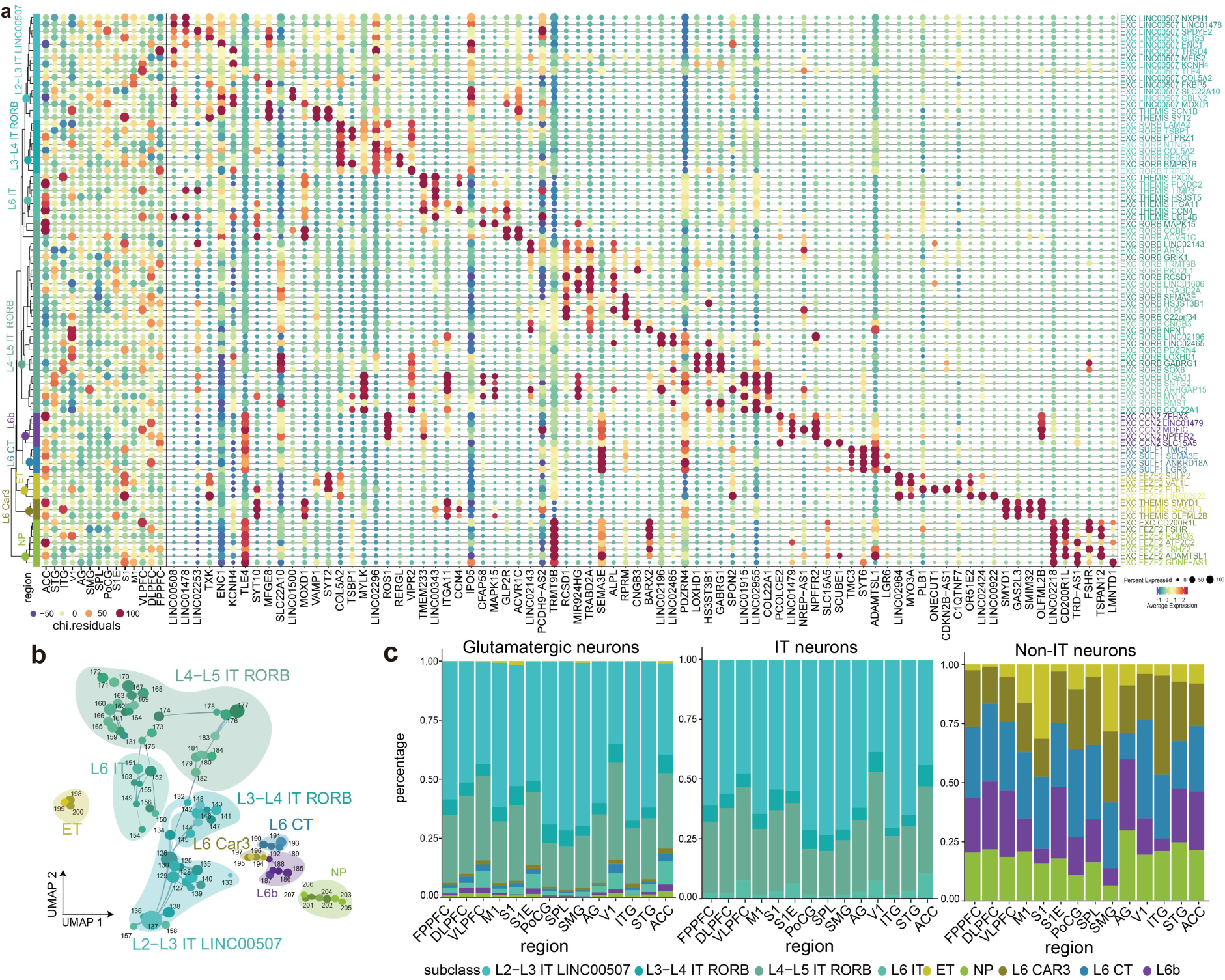
Classification of glutamatergic neurons. **a.** (left) Glutamatergic neurons were classified into 9 subclasses: NP, L6 car3, L6 IT, ET, L6 CT, L6b, L2-L3/IT, L3-L4/IT, and L4-L5/IT. Dot plots below show the distribution preference of cell types across each cortical region. The distribution of preference is calculated by chi-test residuals and further normalized by z-score. (right) The combinational marker for each cluster and their average expression. **b.** Glutamatergic cell-type constellation plot of the global relatedness. Each dot represents a cluster and the link between them denotes the proportion of two clusters sharing which is calculated by KNN. **c.** Bar plot of statistics of glutamatergic subclass at different levels. (Left) Global; (Middle) IT group; (right) other subclasses except for IT.

**Extended Data Fig. 4.**
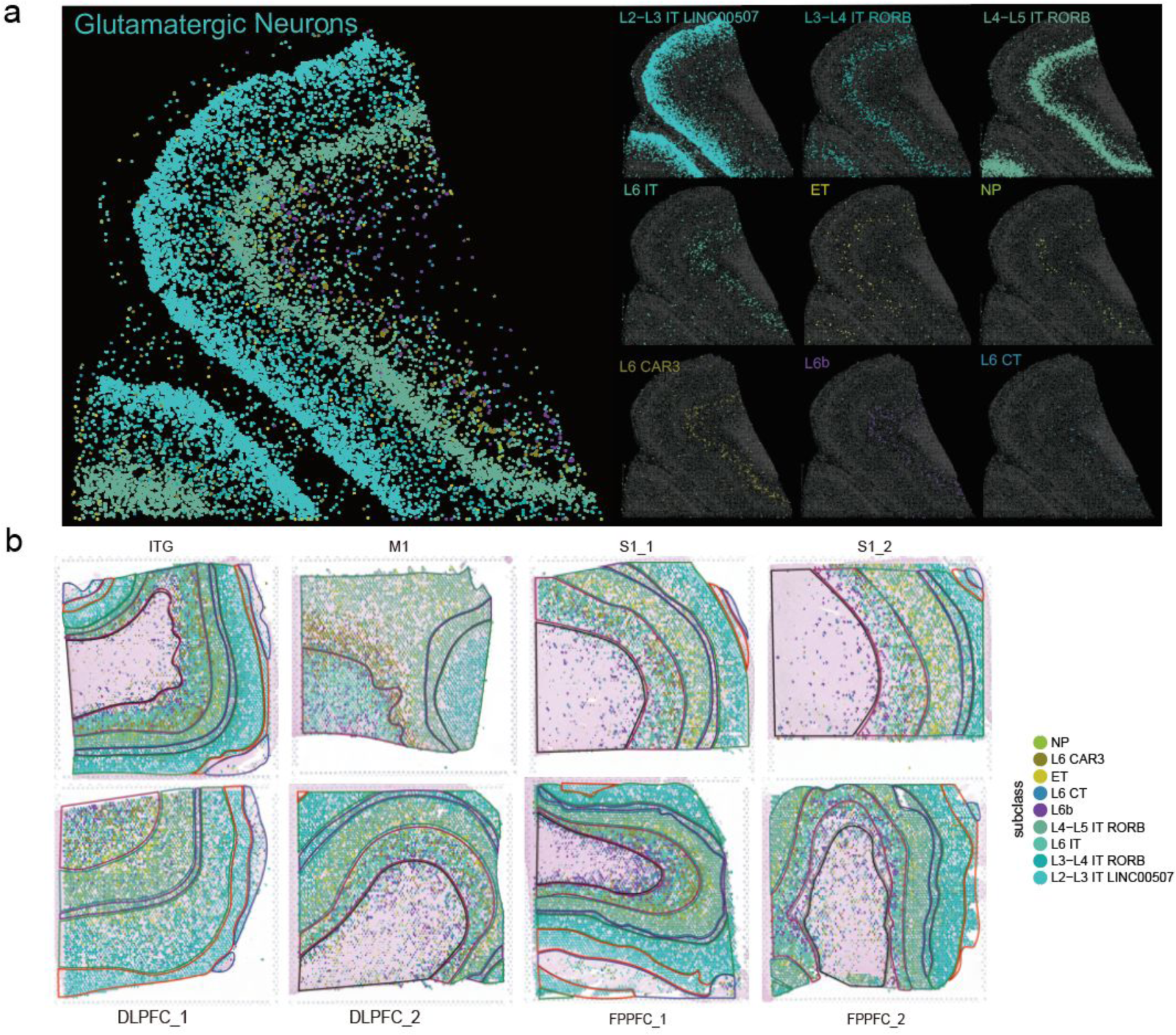
The spatial arrangement of glutamatergic neurons in the slides of the spatial transcriptome. **a.** The cell type annotation for 37.5 μm bins of Stereo-seq section of human DLPFC cortex by snRNA-seq at subclass level with only glutamatergic neurons. Each facet around the main figures denotes a subclass spatial distribution. **b.** Scatterpie showing the proportion of probability of glutamatergic subclasses on different slides. Spots are colored by their subclass identities.

**Extended Data Fig. 5.**
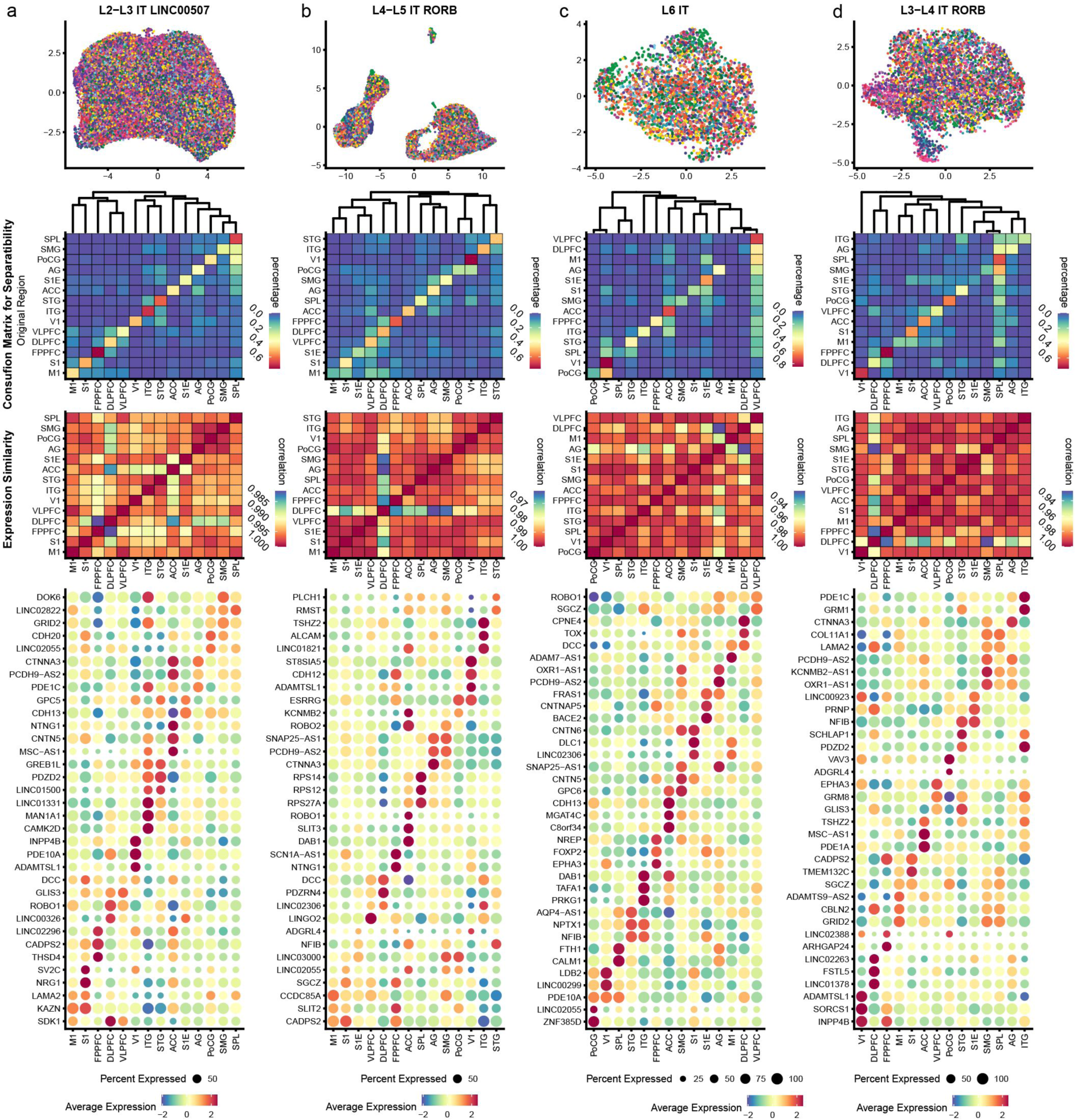
Regional distribution of glutamatergic IT subclasses in the cerebral cortex. **a.** Distribution similarity analysis of L2-L3 IT subclass in cortical regions. From top to bottom, UMAP shows the spatial distribution of all L2-L3 IT cells; the heatmap of the confusion matrix shows the separability of L2-L3 IT cells from distinct cortical regions, and rows and columns correspond to the actual and predicted regional identities of cells, with the rows adding up to 1, with a relevant hierarchy dendrogram on the top; Expression similarity of L2-L3 IT among cortical regions; Heatmap of region-specific marker genes for L2-L3 IT. **b.** Distribution similarity analysis of L4-L5 IT subclass in cortical regions. From top to bottom, the same analysis with **a**. **c.** Distribution similarity analysis of L6 IT subclass in cortical regions. From top to bottom, the same analysis with **a**. **d.** Distribution similarity analysis of L3-L4 IT subclass in cortical regions. From top to bottom, the same analysis with **a**.

**Extended Data Fig. 6.**
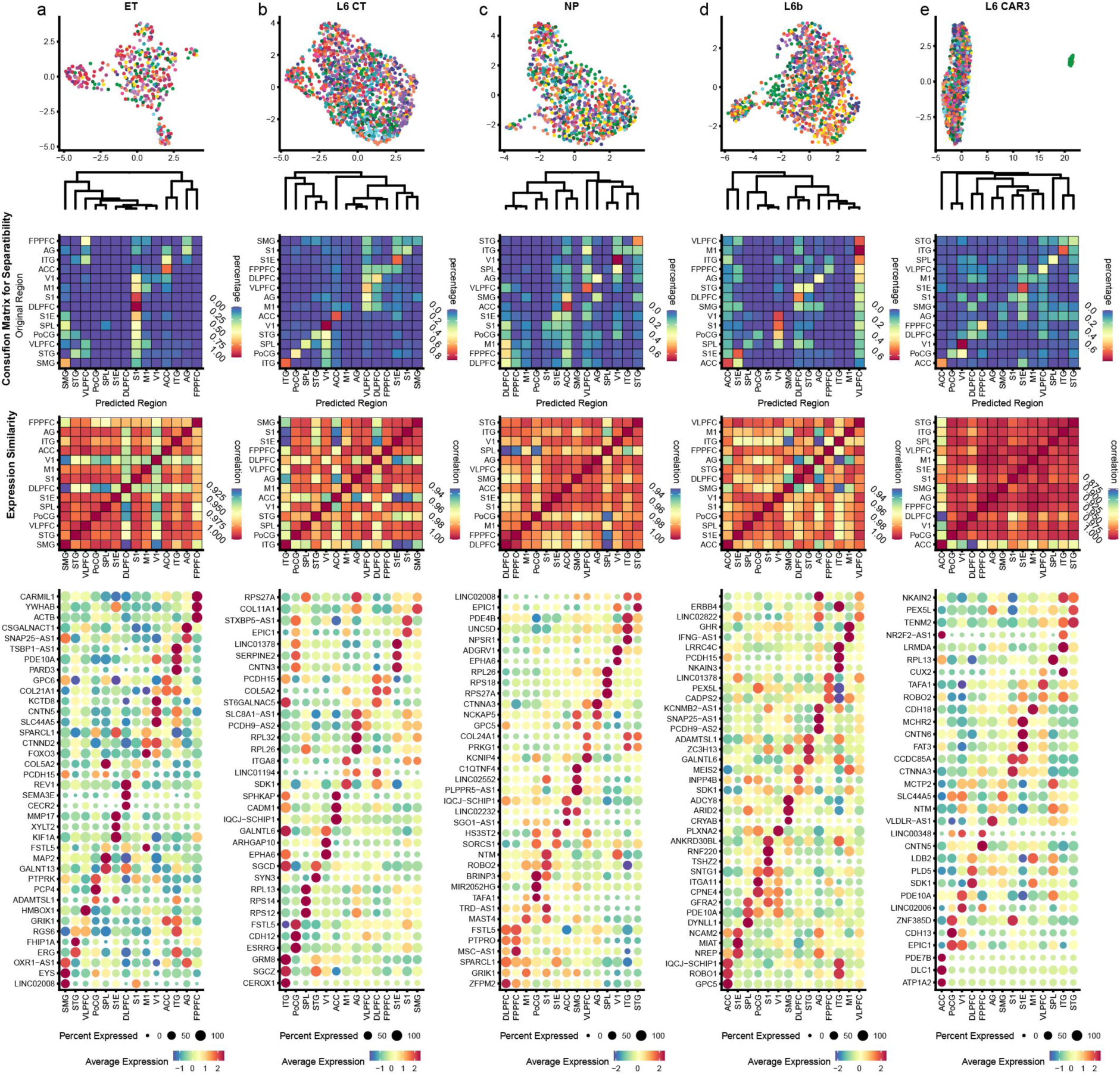
Regional distribution of glutamatergic subclasses except for IT in the cerebral cortex. **a.** Distribution similarity analysis of ET subclass in cortical regions. From top to bottom, the same analysis with Extended Data Fig. 5. **b.** Distribution similarity analysis of L6 CT subclass in cortical regions. **c.** Distribution similarity analysis of NP subclass in cortical regions. **d.** Distribution similarity analysis of L6b subclass in cortical regions. **e.** Distribution similarity analysis of L6 Car3 subclass in cortical regions.

**Extended Data Fig. 7.**
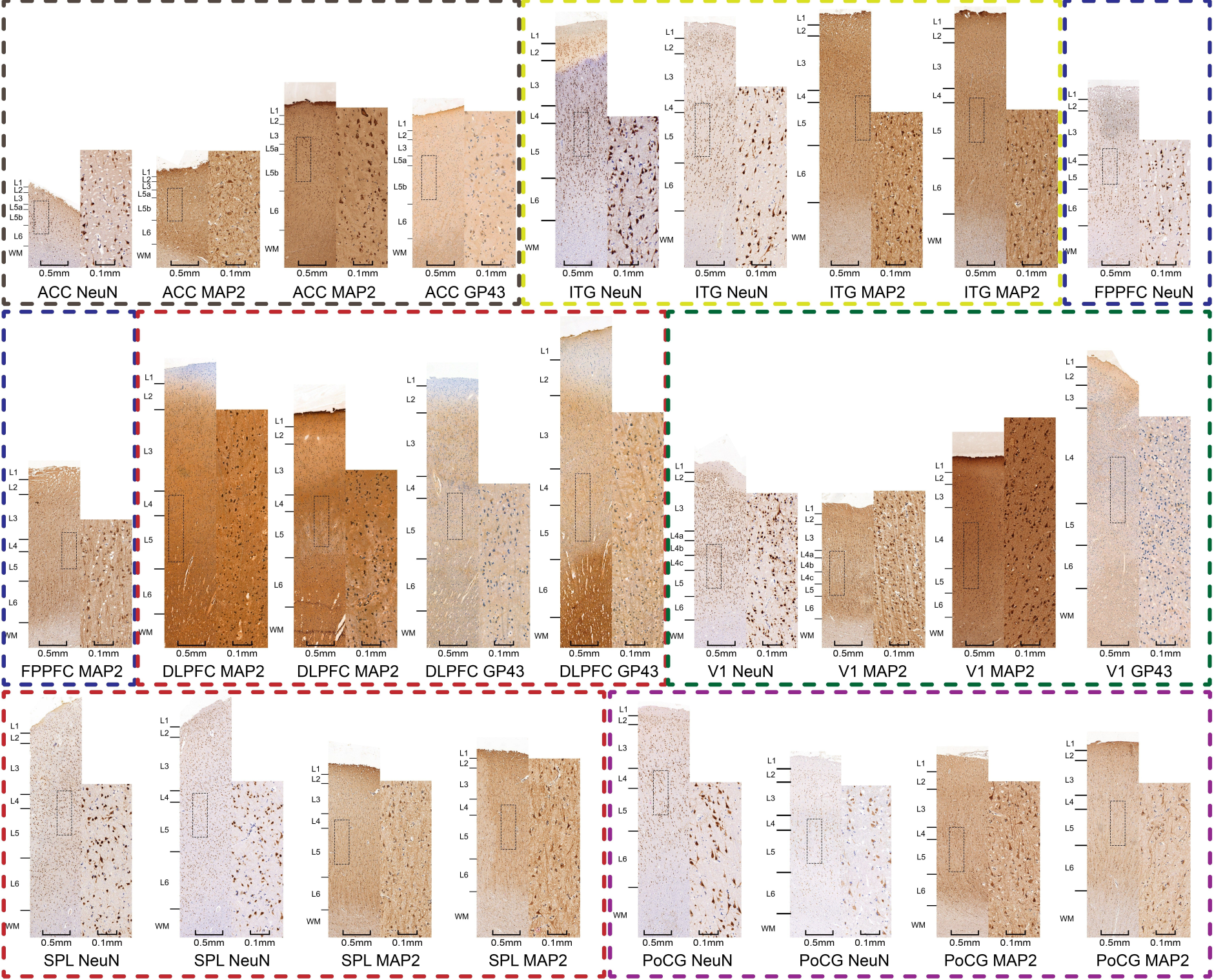
Immunohistochemical staining for many cortical regions.

**Extended Data Fig. 8.**
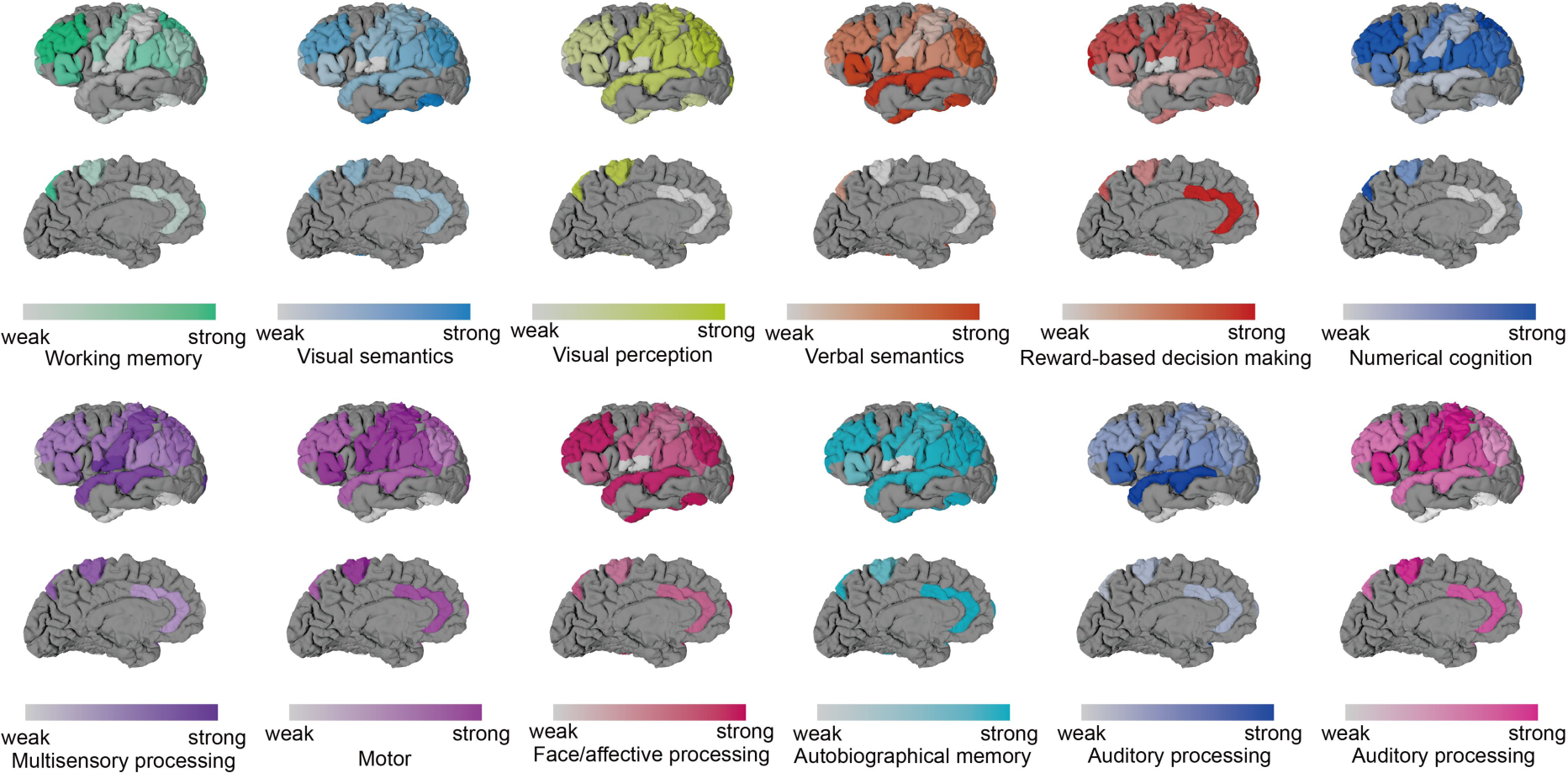
FMRI task correlation with different cortical regions.

**Extended Data Fig. 9.**
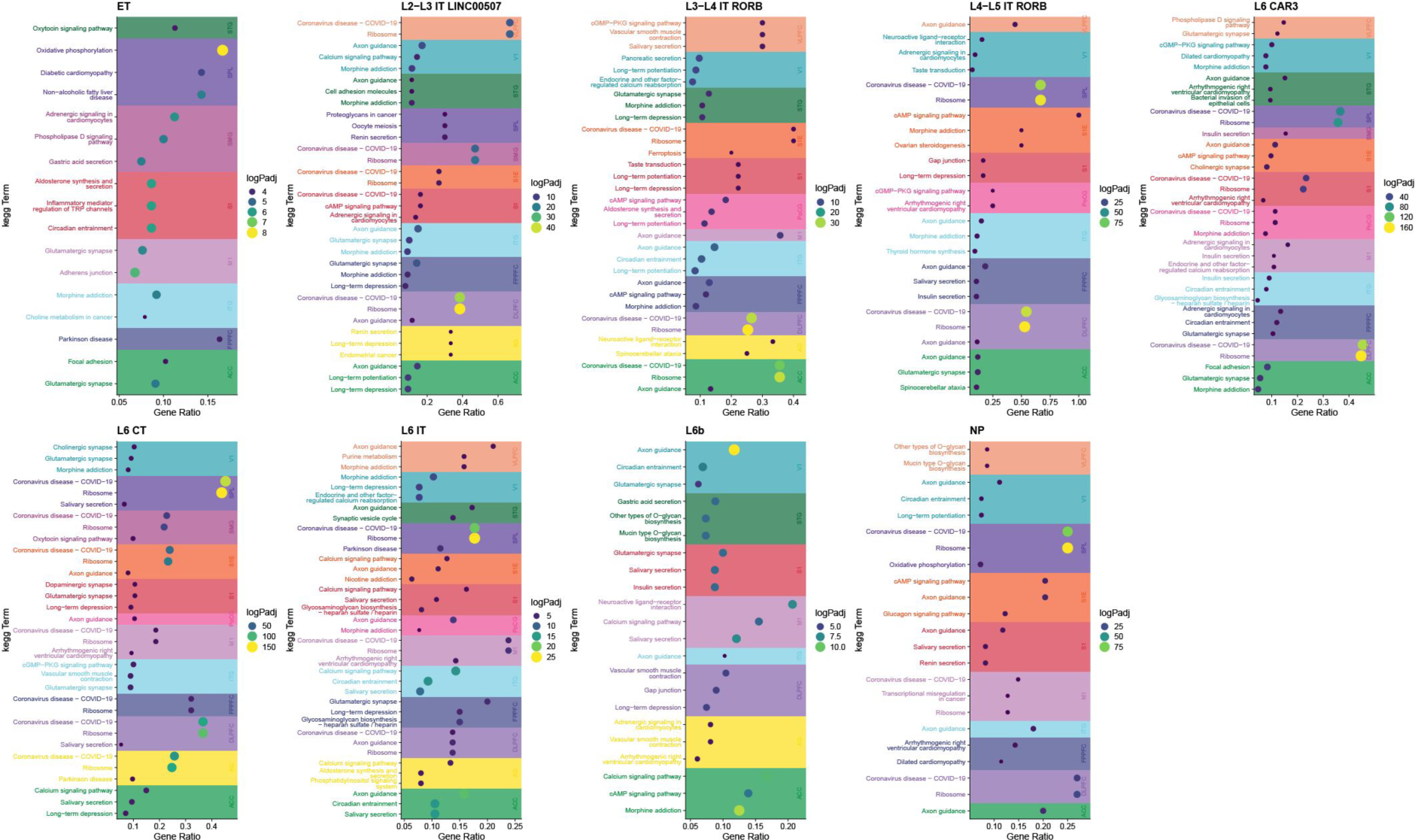
KEGG terms enriched among differentially expressed genes of glutamatergic neurons from each cortical region compared to the rest at the subclass level.

**Extended Data Fig. 10.**
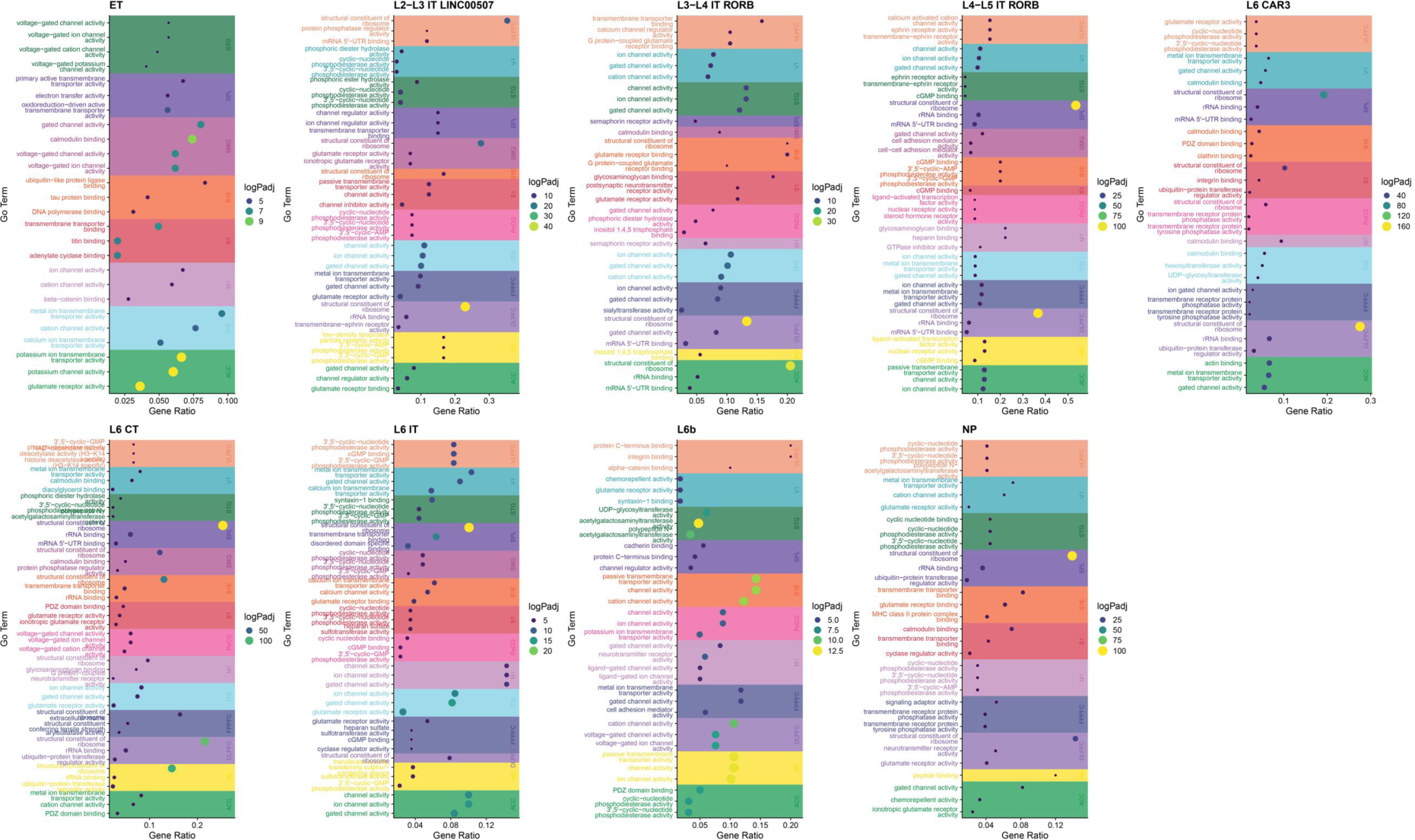
Gene ontology (GO) terms enriched among differentially expressed genes of glutamatergic neurons from each cortical region compared to the rest at the subclass level.

**Extended Data Fig. 11.**
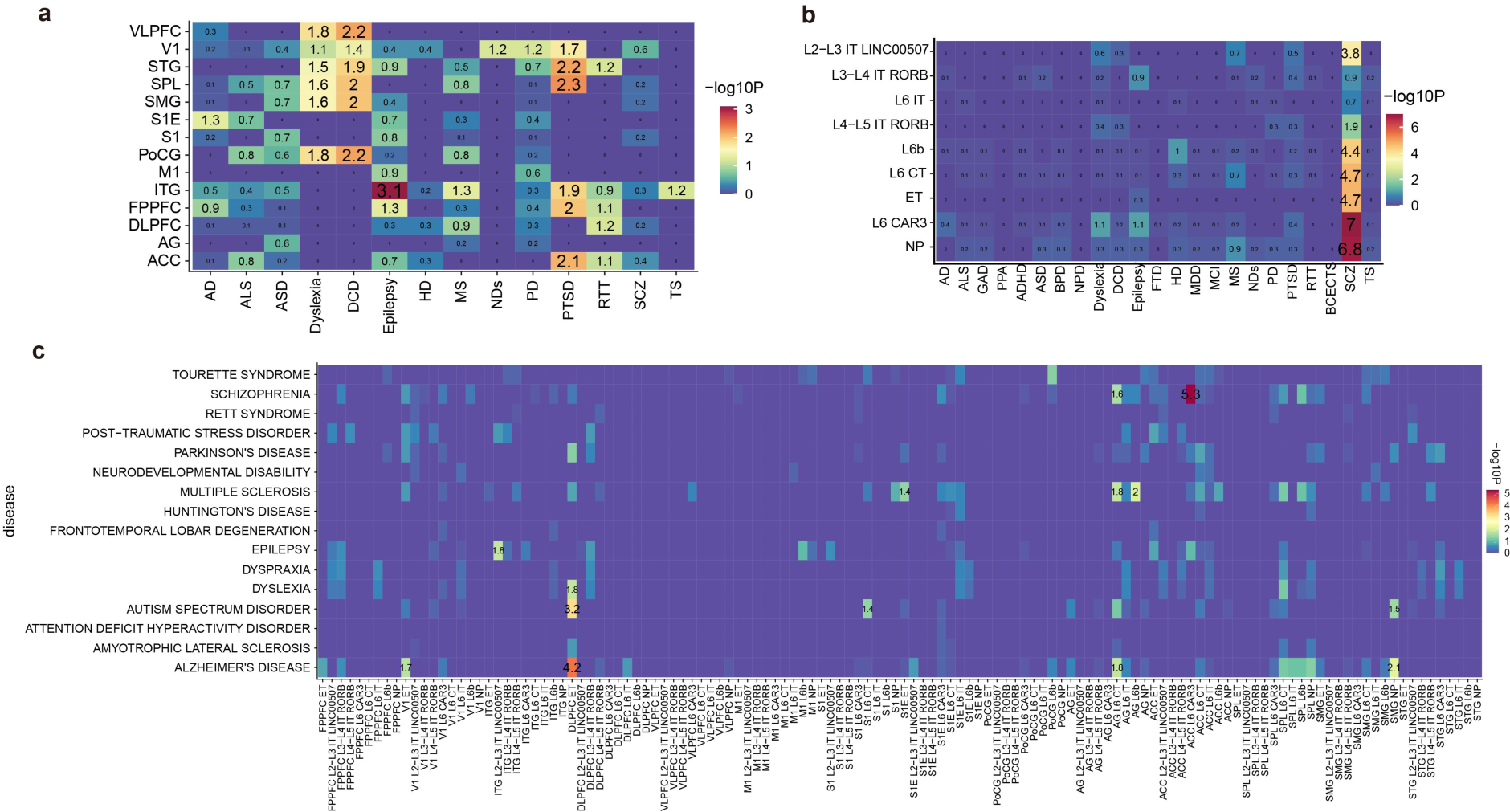
Heatmap of the enrichments of neural disease gene sets for humans among the glutamatergic cell types from snRNA-seq data. Numbers indicate a negative LogP value of the fisher test for enrichment.

**Extended Data Fig. 12.**
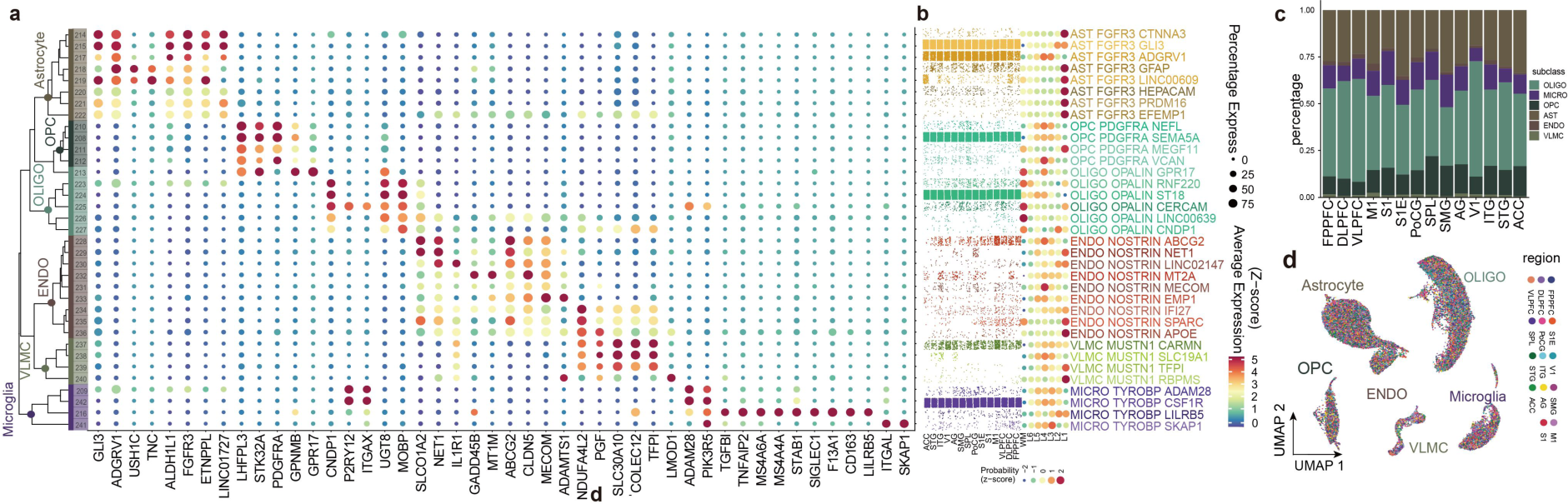
Classification of non-neuronal cells. **a.** Non-neuronal cell-type were divided according to their expressional types into 6 subclasses after snRNA-seq, microglia, VLMC, ENDO, astrocyte, OPC, and oligodendrocyte. Taxonomy of non-neuronal cluster(right). Dot plots showing marker-gene expression distributions across non-neuronal clusters (left). **b.** Scatterplot show regional distribution of non-neuronal cell types, colored by the subclass and dot plot show the laminar specificities calculated by residuals of chi-test of non-neuronal cell types. **c.** Bar plot of statistics of cell number in each subclass. **d.** UMAP representation of non-neuronal subclasses, colored by region.

**Extended Data Fig. 13.**
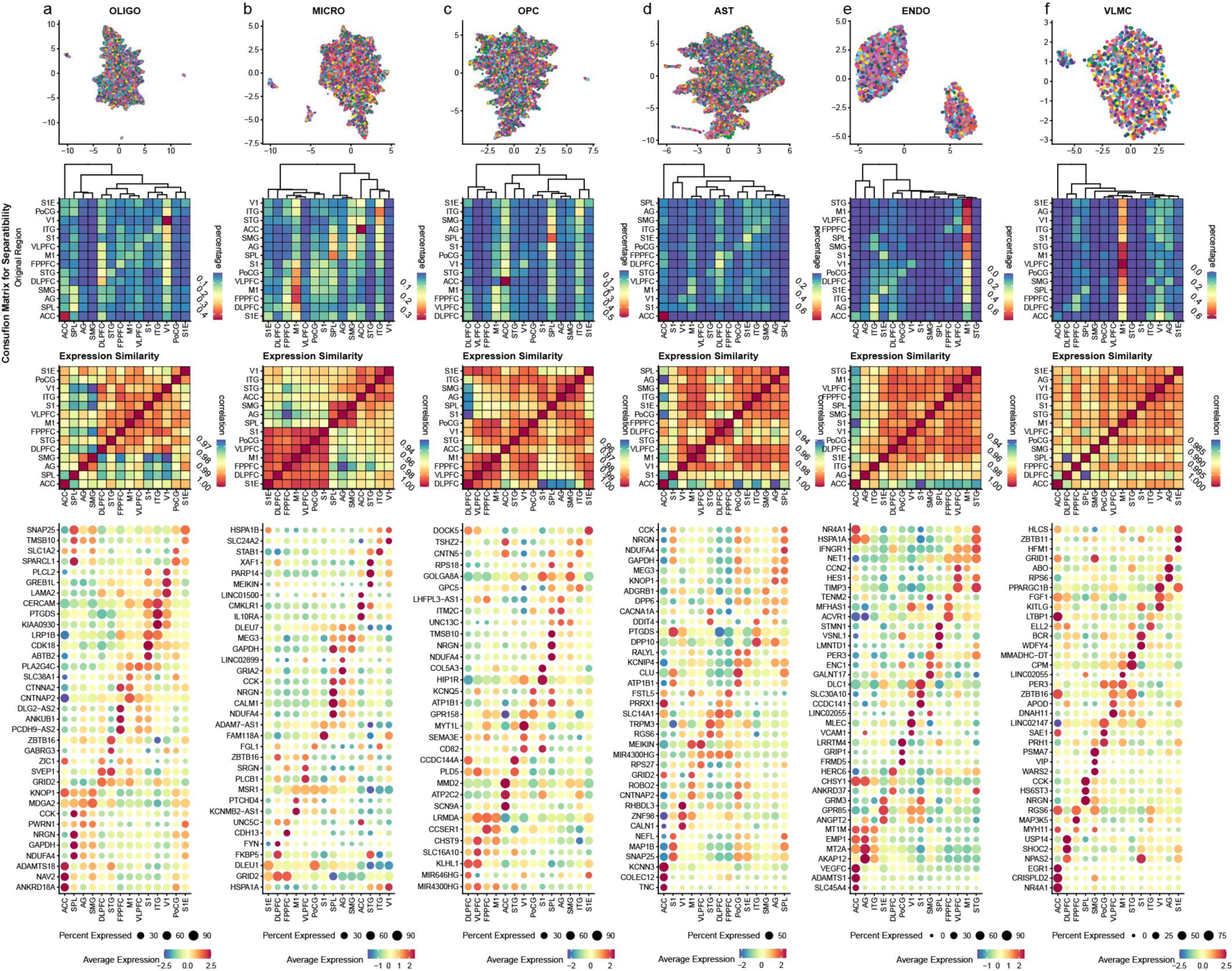
Regional distribution of non-neuronal subclasses in the cerebral cortex. **a.** Distribution similarity analysis of OLIGO subclass in cortical regions. From top to bottom, UMAP shows the spatial distribution of all OLIGO cells; the heatmap of the confusion matrix shows the separability of OLIGO cells from distinct cortical regions, and rows and columns correspond to the actual and predicted regional identities of cells, with the rows adding up to 1, with a relevant hierarchy dendrogram on the top; Expression similarity of OLIGO among cortical regions; Heatmap of region-specific marker genes for OLIGO. **b.** Distribution similarity analysis of microglia subclass in cortical regions. From top to bottom, the same analysis with a. **c.** Distribution similarity analysis of OPC subclass in cortical regions. From top to bottom, the same analysis with a. **d.** Distribution similarity analysis of astrocyte subclass in cortical regions. From top to bottom, the same analysis with a. **e.** Distribution similarity analysis of ENDO subclass in cortical regions. From top to bottom, the same analysis with a. **f.** Distribution similarity analysis of VLMC subclass in cortical regions. From top to bottom, the same analysis with a.

**Extended Data Fig. 14.**
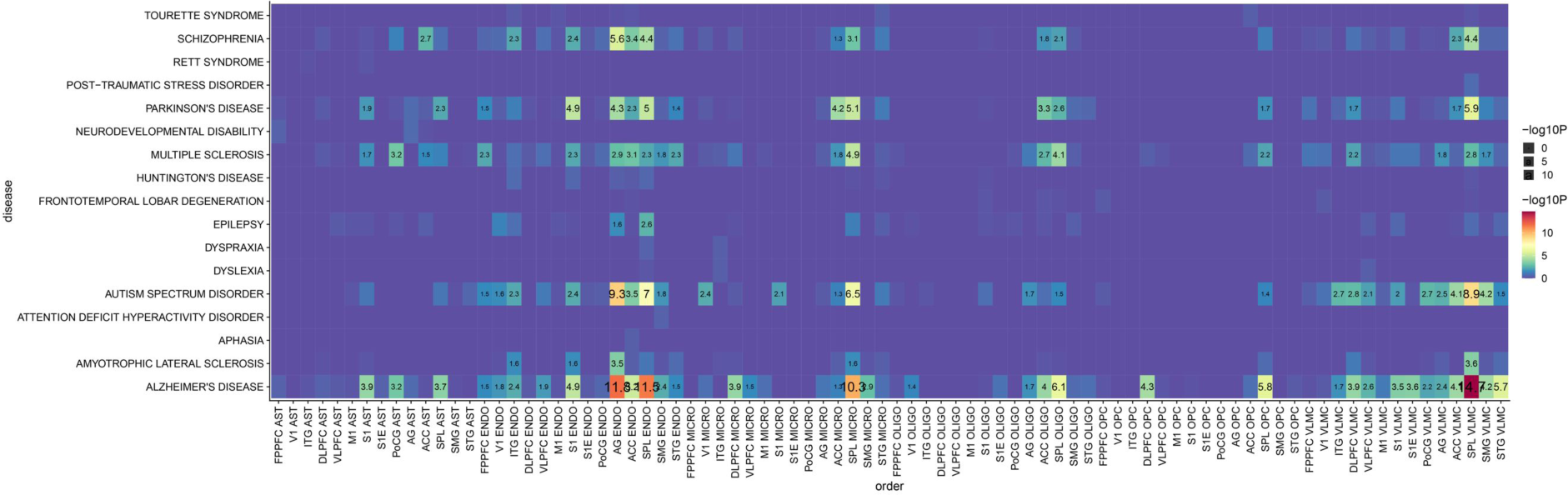
Heatmap of the enrichments of neural disease gene sets for humans among the non-neuronal cell types from snRNA-seq data. Numbers indicate a negative LogP value of the fisher test for enrichment.

**Extended Data Fig. 15.**
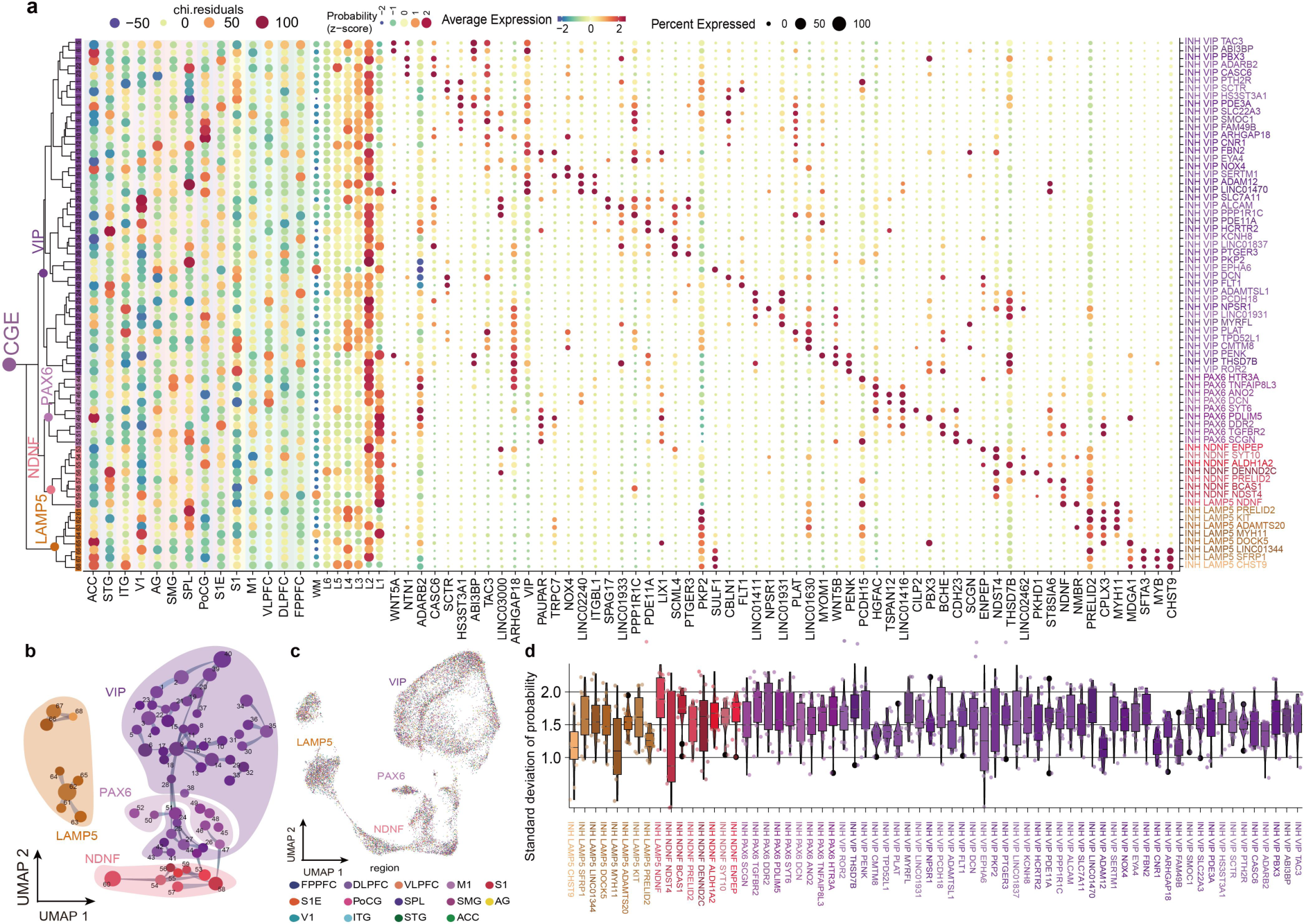
Classification of CGE-derived neurons. **a.** Cells developed from CGE were clustered into 4 subclasses after snRNA-seq, with a total of 35 cell types. Taxonomy of CGE cell types. (left) A dot plot of the distributional chi-test residuals of clusters in each region; (middle) The probabilities of laminar distribution of different clusters; (right) marker-gene expression distributions across CGE-derived cell types. **b.** CGE-derived cell-type constellation plot of the global relatedness. Each dot represents a cluster and the link between them denotes the proportion of two clusters sharing which is calculated by KNN. **c.** UMAP representation of CGE subclasses, colored by region. **d.** Dispersion of laminar distribution by CGE-derived cell types. Each point denotes the standard deviation of the probability of each cell type on each slide. A low value indicates a dense density of distribution.

**Extended Data Fig. 16.**
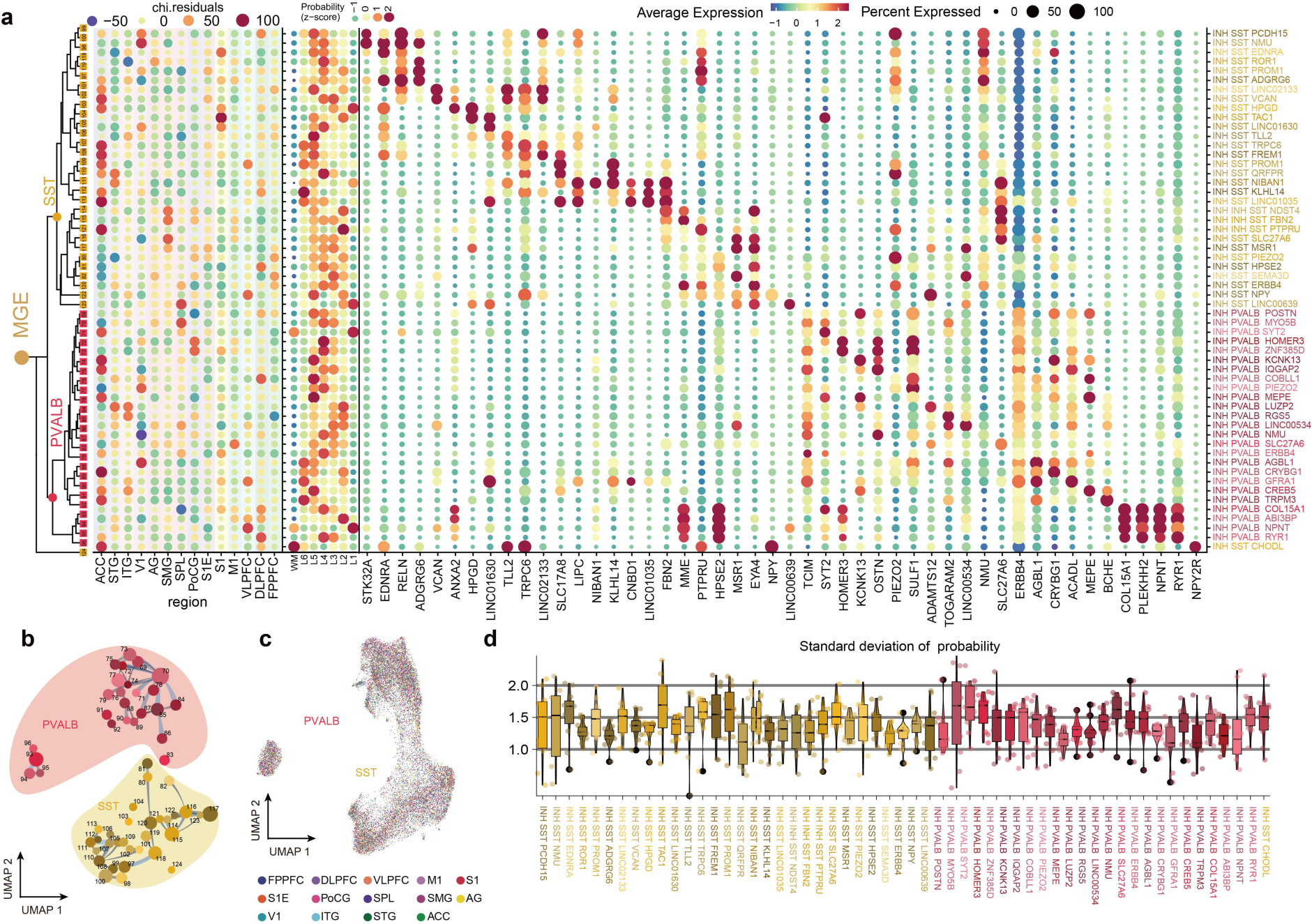
Classification of MGE-derived neurons. **a.** Taxonomy of MGE-derived neurons. (left) A dot plot of the distributional chi-test residuals of MGE-derived cell types in each region; (middle) The probabilities of laminar distribution of different clusters; (right) marker-gene expression distributions across MGE-derived cell types. **b.** MGE-derived cell-type constellation plot of the global relatedness. The same with Extended Data Fig. 15b. **c.** UMAP representation of MGE subclasses, colored by region. The same with Extended Data Fig. 15c. **d.** Dispersion of laminar distribution by MGE-derived cell types. The same with Extended Data Fig. 15d.

**Extended Data Fig. 17.**
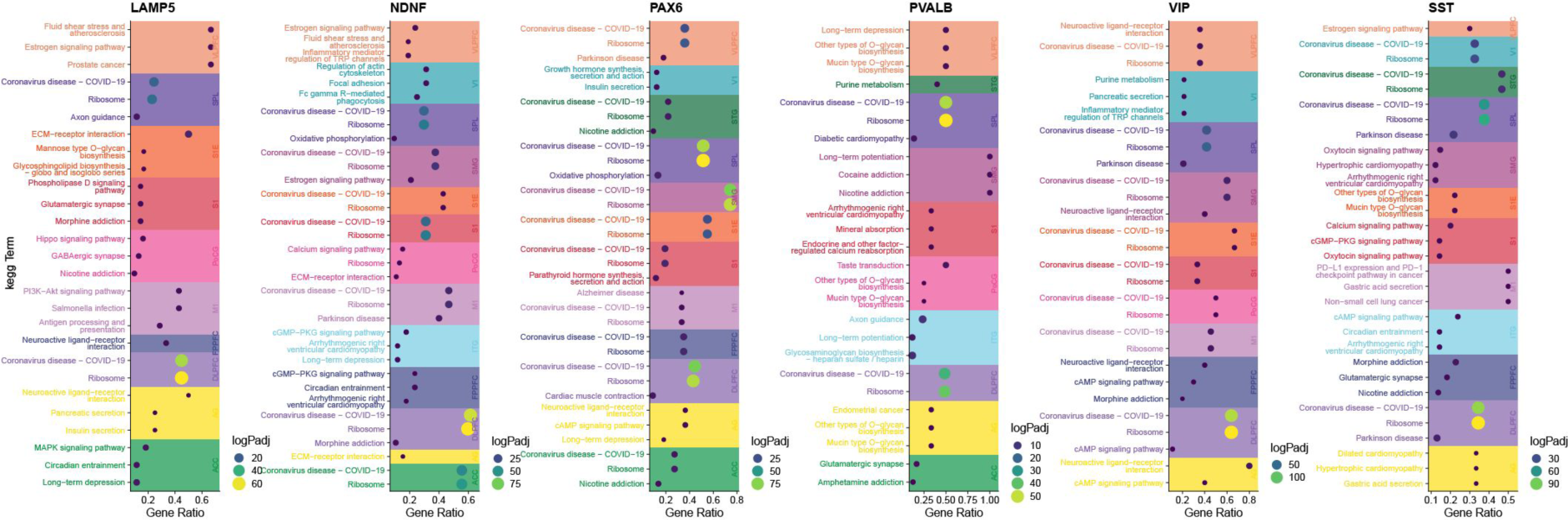
KEGG terms enriched among differentially expressed genes of GABAergic neurons from each region compared to the rest at the subclass level.

**Extended Data Fig. 18.**
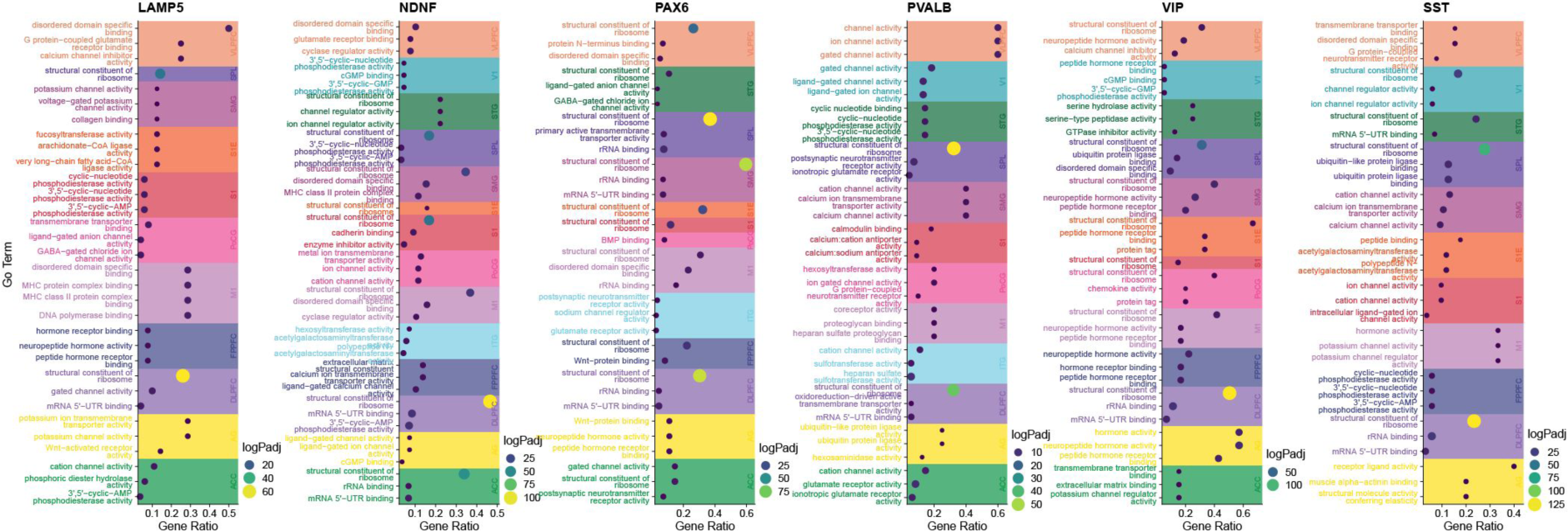
GO terms enriched among differentially expressed genes of GABAergic neurons from each region compared to the rest at the subclass level.

**Extended Data Fig. 19.**
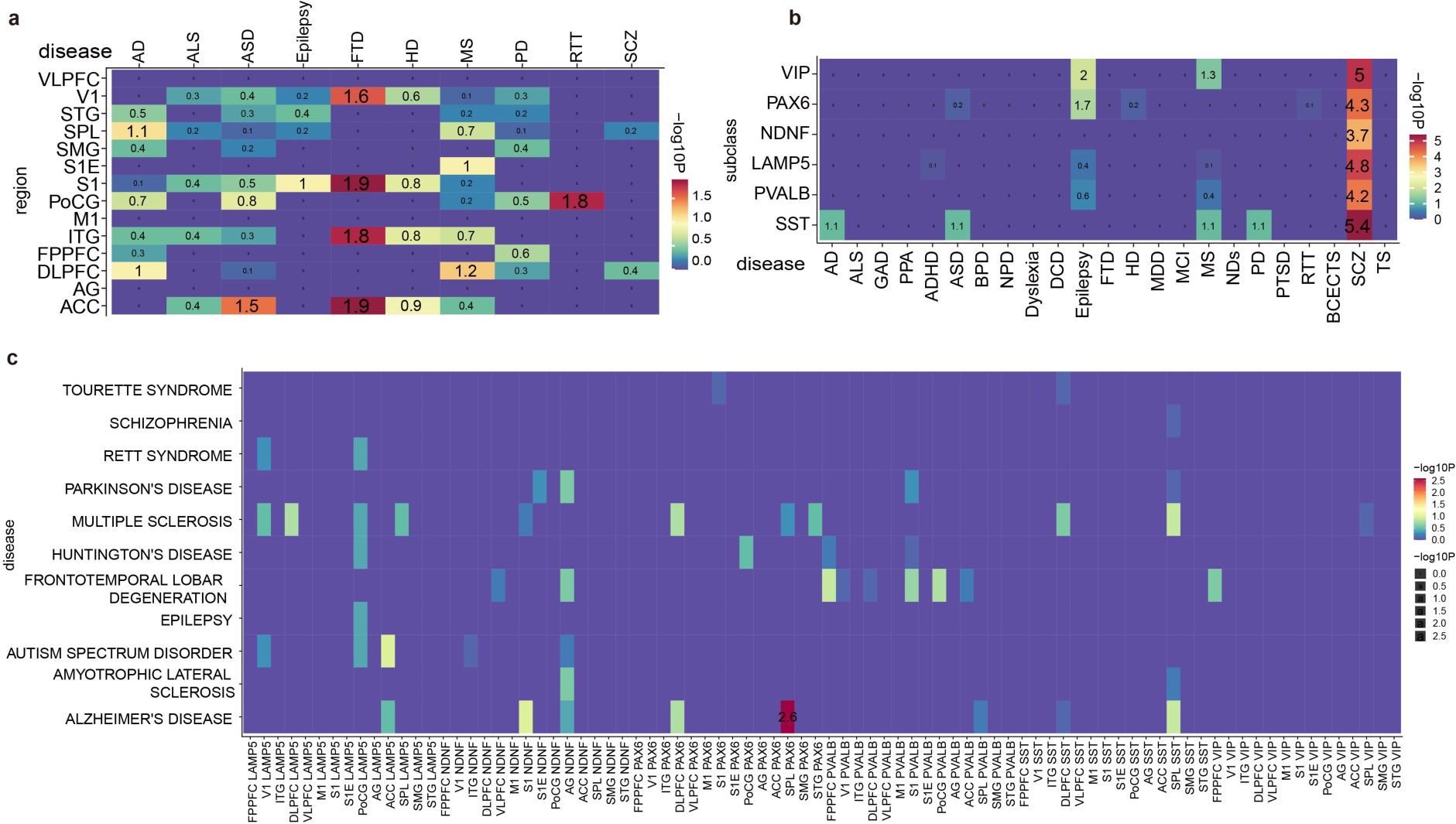
Heatmap of the enrichments of neural disease gene sets for humans among the GABAergic cell types from snRNA-seq data. Numbers indicate a negative LogP value of the fisher test for enrichment.

## Supplemental tables

**Supplementary Table 1. Sample information and mRNA alignment.**

Sample information, and single-cell and spatial transcriptomic alignment data.

**Supplementary Table 2. Laminar markers and data of cell-type deconvolution.**

Markers for each cortical layer, and data of deconvolution of each cell type across 14 cortical regions.

**Supplementary Table 3. Regional expression difference among subclass, pathway enrichment, and disease susceptibility.**

Markers of different cell types in different regions, and their enrichment in GO, Kegg, and disease gene sets.

**Supplementary Table 4. Data of reclustering on spots from WM and layer 6.**

Statistics of clustering results on WM and layer 6, and differential gene expression and specific cell types in layer 6b.

